# Highly sensitive quantification of carbon monoxide (CO) *in vivo* reveals a protective role of circulating hemoglobin in CO intoxication

**DOI:** 10.1101/2020.10.12.336735

**Authors:** Qiyue Mao, Akira T. Kawaguchi, Shun Mizobata, Roberto Motterlini, Roberta Foresti, Hiroaki Kitagishi

## Abstract

Carbon monoxide (CO) is a gaseous molecule known as the silent killer. It is widely believed that an increase in blood carboxyhemoglobin (CO-Hb) is the best biomarker to define CO intoxication, neglecting the important fact that CO accumulation in tissues is the most likely direct cause of mortality. There is no reliable method other than gas chromatography to accurately determine CO content in tissues. Here we report the properties and usage of hemoCD1, a synthetic supramolecular compound composed of an iron(II)porphyrin and a cyclodextrin dimer, as an accessible reagent for a simple colorimetric assay to quantify CO in biological samples. The assay was validated in various organ tissues collected from rats under normal conditions and after exposure to CO by inhalation. The kinetic profile of CO in blood and tissues after CO treatment suggested that CO accumulation in tissues is prevented by circulating Hb, revealing a protective role of Hb in CO intoxication. Furthermore, hemoCD1 was used *in vivo* as a CO removal agent, showing that it acts as effective adjuvant to O_2_ ventilation to eliminate residual CO accumulated in organs, including the brain. These findings open new therapeutic perspectives to counteract the toxicity associated with CO poisoning.

## Introduction

Carbon monoxide (CO) is a diatomic gaseous molecule produced by incomplete combustion of carbon-based materials and is mostly known for its toxic properties when inhaled in high amounts. CO intoxication, which frequently occurs in suicides, during fires, and other accidents, is the most common type of chemical poisoning with an increased risk of death (*1*). In spite of its notorious toxicity, CO is relatively unreactive compared to molecular oxygen (O_2_), nitric oxide (NO), hydrogen sulfide (H_2_S), and reactive oxygen species (ROS) such as superoxide (O_2_^−^), hydroxyl radical (•OH), hydrogen peroxide (H_2_O_2_), and singlet oxygen CO_2_). Although other small molecules show various reactivities towards proteins, lipids, and nucleic acids, the only recognized reaction of CO in the biological system is the binding to low valent metal ions, mostly ferrous heme (*2–5*). Due to a specific interorbital interaction, called π-backbonding or π-backdonation, the carbon atom of CO forms a particularly strong bond with low valent metal ions such as iron(II)heme with a high stability and high binding affinity (*6,7*). Formation of the stable heme-CO complex *in vivo* is the cause of CO toxicity; once CO is inhaled, it strongly binds to heme in hemoproteins that are indispensable for carrying or consuming O_2_, such as hemoglobin (Hb), myoglobin (Mb) and cytochrome *c* oxidase (C*c*O) among others, leading to their dysfunction (*8,9*). Levels of CO-bound Hb (CO-Hb) in blood are considered an important biomarker to assess CO poisoning (*1,10,11*). CO-Hb below 10% normally circulates in blood due to endogenously produced CO or CO from cigarette smoking, with no apparent clinical symptoms. When CO-Hb reaches levels between 20 to 40% due to CO gas inhalation, symptoms such as dizziness, headache, and nausea begin to appear. CO-Hb values exceeding 40% increase the risk of death. Since the binding of CO to heme is thermodynamically controlled and thus reversible, CO-Hb levels can be reduced by excess O_2_. O_2_ ventilation is thus a current major approach for treating CO intoxication (*12,13*).

A unique study conducted in dogs and reported in 1975 (*14*) provides an important insight on the mechanism of CO poisoning. When dogs received CO gas by inhalation, they all died rapidly within 1 h with CO-Hb levels of 54 to 90%. In contrast, if the animals were first bled to reduce blood volume and then transfused with red blood cells (RBC) saturated with CO, all dogs survived despite reaching CO-Hb levels of 60% (*14*). These findings strongly indicate that inhalation of gaseous CO is highly toxic whereas CO-Hb is not, ultimately challenging the commonly accepted notion that the percentage of CO-Hb is the appropriate marker to establish the toxicity of CO inhalation. Indeed, patients can die of CO poisoning even with CO-Hb saturation levels below 30% (*13,15*), indicating that crucial toxicity derives from gaseous CO diffusing into organs/tissues, directly compromising the activities of key hemoproteins such as *CcO* that requires O_2_ to produce energy (*8*). From the viewpoints of medical diagnosis and toxicology, knowing how much CO is accumulated in organs/tissues in subjects affected by CO intoxication would be fundamentally important.

Detection and quantification of CO contained in biological samples have been a long-standing challenge. In medical practice, CO-Hb in blood is measured by oximeters, which rely on a colorimetric assay to diagnose CO intoxication (*10*). In addition, gas chromatography (GC) techniques are the main methodology used to detect CO contained in tissues, as reported by the laboratories of Coburn and Vreman to determine endogenous CO distributed in mice (*16*) and human tissues (*17,18*). GC is considered to be well established as a CO quantifying method (*10,19*). However, as we investigated and discuss in this paper, the headspace analysis by GC tends to underestimate the amount of CO, highlighting that a convenient and reliable method for measuring CO contained in organs/tissues is needed.

Biomimetic chemistry for hemoproteins started in the 1970s. An epoch-making model for O_2_-binding hemoproteins is a picket-fence porphyrin synthesized by Collman and co-workers (*20–22*). After their discovery, many synthetic analogs were synthesized to demonstrate the reversible O_2_/CO bindings in anhydrous organic solvents (*21,22*). The weakness of these synthetic model systems was that even a trace amount of water contaminating the solvent was unacceptable due to a water-catalyzed autoxidation of the iron(II) to iron(III)porphyrins (*23,24*). Because of the difficulty in preparing a hydrophobic heme pocket similar to native hemoproteins, aqueous biomimetic compounds have been scarce (*25,26*). Nevertheless, our laboratory has succeeded in preparing hemoprotein models working in 100% aqueous solutions (*27–33*). In our model system, the iron(II)porphyrin is embedded in hydrophobic per-*O*-methyl-β-cyclodextrin (CD) cavities, thus avoiding a water-catalyzed autoxidation even in 100% water at ambient temperatures. As a result, stable and reversible O_2_ and CO bindings similar to native Hb and Mb have been achieved.

Among the hemoprotein model complexes synthesized in our laboratory, hemoCD1 (Fig. 1A) composed of 5,10,15,20-tetrakis(4-sulfonatophenyl)porphinatoiron(II) (Fe^II^TPPS) and a per-*O*-methyl-β-cyclodextrin dimer with a pyridine linker (Py3CD), showed the highest CO binding affinity (*P*_1/2_^CO^ = 1.5 × 10^−5^ Torr, *K*d = 19.2 pM in phosphate buffer at pH 7 and 25°C) (*28*). This is approximately 100 times higher than that of Hb in R-state (Hb-R), whereas the O_2_ binding affinity of hemoCD1 is moderate (*P*_1/2_^O2^ = 10 Torr, *K*_d_ = 17 μM in phosphate buffer at pH 7 and 25°C) (*28,34*), close to that of Hb in the T-state (Hb-T). To the best of our knowledge, the CO binding affinity of hemoCD1 is the highest among the reported native CO-binding proteins. The administration of hemoCD1 to animals (rats and mice) did not show any significant side effect except for endogenous CO scavenging (*35–38*). Most of the administered hemoCD1 is excreted in the urine as CO-hemoCD1 complex within one hour. Therefore, we have focused on studying hemoCD1 as a CO scavenging agent *in vivo* and recently described the effects of endogenous CO-depletion on heme oxygenase expression (*35*), ROS production (*39*), and circadian clock gene expressions (*38*). For years we have recognized that hemoCD1 not only is a powerful tool for elucidating the biological role of endogenous CO but could also be useful for detecting and removing excessive CO in the living organisms.

**Fig. 1.**
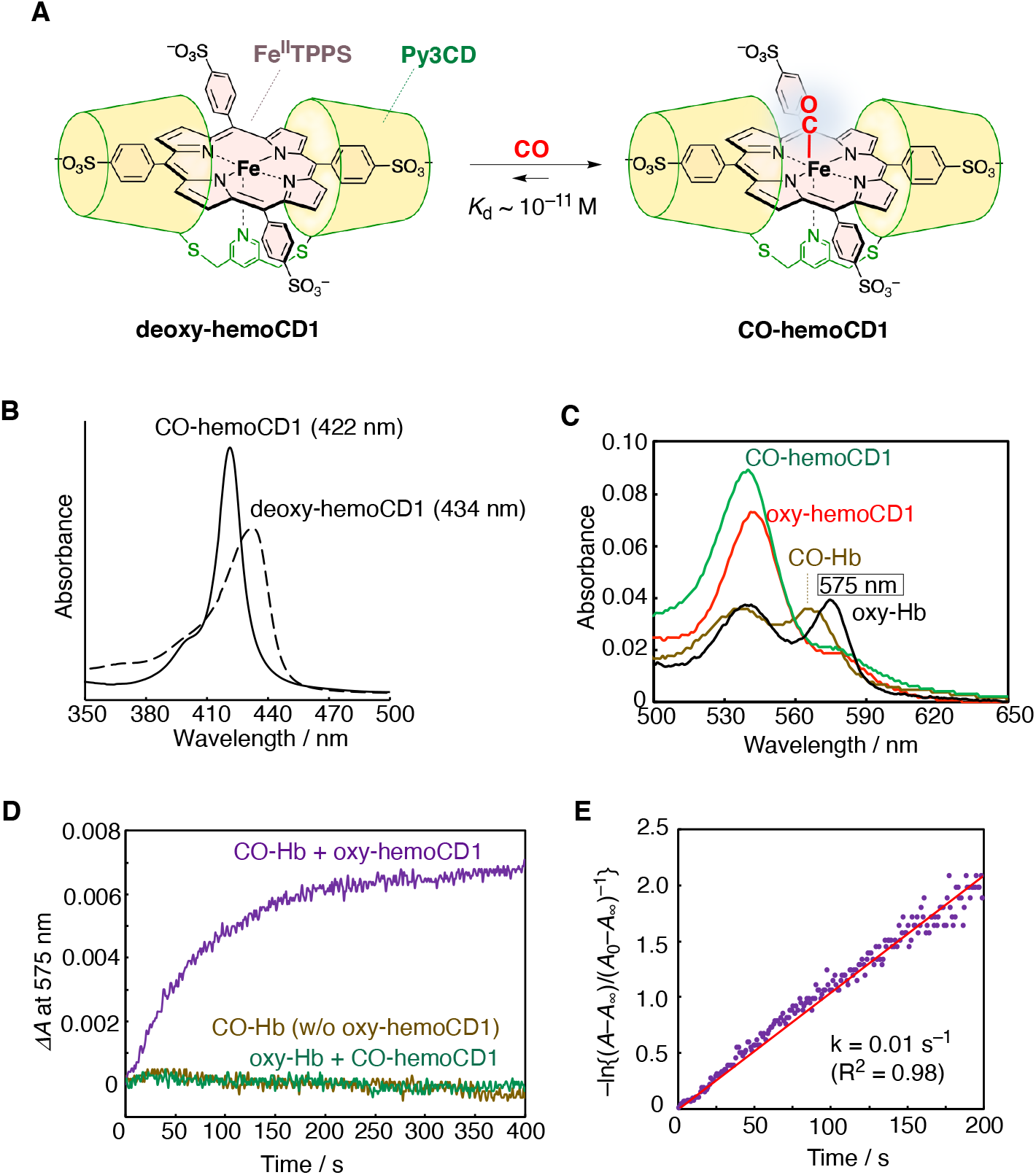
HemoCD1, a CO detecting agent. (**A**) HemoCD1 is composed of 5,10,15,20-tetrakis(4-sulfonatophenyl)porphinatoiron(II) (Fe^II^TPPS) and a per-*O*-methyl-β-cyclodextrin dimer having a pyridine linker (Py3CD). The structure of deoxy-hemoCD1 and CO-hemoCD1 complexes is shown. (**B**) UV-vis spectra of hemoCD1 showing the Soret bands typical of deoxy-hemoCD1 (434 nm) and CO-hemoCD1 (422 nm) in PBS at pH 7.4 and 25°C. (**C–E**) Competition between hemoCD1 and Hb for CO binding. (**C**) UV-vis spectra of oxy-, and CO-hemoCD1 (5.0 μM each), and oxy-(575 nm absorbance peak) and CO-Hb (2.4 μM each). (**D**) Time-course for changes in absorbance at 575 nm, indicative of formation of oxy-Hb, after mixing stock solutions of oxy-hemoCD1 (0.75 mM, 20 μL in air-saturated PBS) and CO-Hb (0.72 mM, 10 μL in CO-saturated PBS) in air-saturated PBS (3 mL) at pH 7.4 and 25°C. Controls are represented by solutions of CO-Hb without (w/o) oxy-hemoCD1, and oxy-Hb mixed with CO-hemoCD1. (**E**) First-order rate plot for changes in absorbance at 575 nm over time. The linear regression analysis gave a rate constant of *k* = 0.01 s^−1^.

In this paper, we established a new method for CO detection and quantification using hemoCD1 in rat tissues. The binding of CO to hemoCD1 is determined by a simple colorimetric assay using absorbances at three different wavelengths (422, 427, and 434 nm) and was evaluated in rats exposed to CO gas compared to untreated animals. Importantly, kinetic studies on bio-distribution of CO in tissues after CO gas inhalation suggest a protective effect of circulating Hb in CO intoxication. Finally, we describe the potential use of hemoCD1 as an antidote for CO poisoning in mammals.

## Results

### HemoCD1 as a suitable and effective CO scavenger

The structure of hemoCD1 is shown in Fig. 1. In the presence of Na_2_S_2_O_4_, the oxidized form of hemoCD1 (met-hemoCD1) is converted to deoxy-hemoCD1 (Fig. 1A), with a typical absorption band at 434 nm (Fig. 1B). Upon addition of CO gas, deoxy-hemoCD1 is smoothly converted into its CO-bound form, CO-hemoCD1 (Fig. 1A), which has a distinct absorption band at 422 nm (Fig. 1B).

Once formed, CO-hemoCD1 is quite stable. In fact, the absorption band of CO-hemoCD1 was unchanged when the solution was bubbled with pure O_2_ for 5 min (Fig. S1). On the other hand, CO-Hb in solution was easily converted to its O_2_-bound form following addition of O_2_ (Fig. S1). Once formed, CO-hemoCD1 could not be reconverted to its deoxy form despite continuous N2 bubbling for 60 min, while CO-Hb returned to its deoxy-form after N2 treatment (Fig. S2). In addition, CO-hemoCD1 remained stable even after the solvent was almost evaporated under high vacuum (<10 Torr) over 2 h. The redissolved complex still showed the distinct and typical spectrum of CO-hemoCD1 (Fig. S3). In contrast, CO-Hb significantly decomposed after the same treatment (Fig. S3). CO-hemoCD1 was also resistant against excess H_2_O_2_, whereas CO-Hb was degraded by H_2_O_2_ under the same conditions (Fig. S4). CO-hemoCD1 is also stable over 24 h in the presence of biocomponents from cell lysates (Fig. S5). These data demonstrate the high stability of the CO-hemoCD1 complex and emphasize that, once bound to hemoCD1, CO hardly dissociates even under these extreme conditions.

To demonstrate the CO-scavenging ability of hemoCD1, CO transfer from CO-Hb to oxy-hemoCD1 (an O_2_-adduct of hemoCD1) was investigated. The spectra of each component are shown in Fig. 1C and we note the peak in absorbance at 575 nm characteristic of oxy-Hb. After mixing the solutions of oxy-hemoCD1 (0.75 mM, 20 μL in air-saturated buffer) and CO-Hb (0.72 mM, 10 μL in CO-saturated buffer where [CO] = 0.96 mM (*40*)) in 3 mL air-saturated buffer, a time-dependent increase in the absorbance at 575 nm was observed due to the conversion of CO-Hb to oxy-Hb (Fig. 1D). The two controls, namely 1) the CO-Hb solution without (w/o) oxy-hemoCD1, and 2) the mixed solution of oxy-Hb with CO-hemoCD1, did not reveal any change in absorbance, as shown in Fig. 1D. The first-order CO transfer rate constant was determined (Fig. 1E), which is consistent with the dissociation rate constant (*k*off^CO^ = 0.01 s^−1^) reported for CO-Hb (*41*). In the CO transfer reaction, dissociation of CO from CO-Hb is the rate-limiting step, as the on rate for CO to hemoCD1 is much faster (*k*_on_^CO^ = 4.7 × 10^7^ M^−1^s^−1^) (*28*). After ultrafiltration of the mixed solution using a filter of molecular weight cut off = 30,000 Da, the hemoCD1 and Hb components were separated. The filtrate contained CO-hemoCD1 whereas the residue contained oxy-Hb (Fig. S6).

Concerning NO, we have already shown that hemoCD1 is unable to bind NO in the presence of Hb, due to its lower NO binding affinity compared to Hb (*39*). In addition, CO-hemoCD1 was not decomposed by the addition of NO, whereas the CO in CO-Hb was replaced by NO leading to formation of met-Hb (Fig. S7). The stability of CO-hemoCD1 even in the presence of NO is an additional advantage for a selective and accurate detection of CO *in vivo*. As for H_2_S, we previously reported that the coordination strength of H_2_S to iron(II) in hemoCD1 is relatively weak, and the SH^−^ ligand is easily replaceable with CO in the hemoCD1 analog (*42*).

The intrinsic toxicity of the CD cavity, which was suggested in a recent article (*8*), was not observed because the CD cavity is occupied by Fe^II^TPPS in hemoCD1 (Fig. S8). We have already reported that the hemoCD1 components have no effect on hemodynamic parameters such as heart rate, blood pressure, and plasma components (*35,43*). It has been demonstrated that biological systems do not recognize *meso*-tetraarylporphinatoiron such as FeTPPS in hemoCD1 as a heme cofactor (*44,45*) and thus hemoCD1 cannot be metabolized by heme oxygenase. Moreover, we demonstrated that injection of oxy-hemoCD1 in rats results in excretion of CO in the urine (*35*) and that the ferric iron(III) form the hemoCD derivatives function as cyanide-antidotes (*32,46,47*). Altogether, our data support the idea that hemoCD1 is a safe and effective CO scavenger.

### Development of a new and sensitive CO quantification method using hemoCD1

Absorbances of the Soret bands of deoxy-hemoCD1 (434 nm) and CO-hemoCD1 (422 nm) were used for CO quantification (Fig. 1B). When aqueous CO solutions with different CO contents were mixed with deoxy-hemoCD1, the Soret band clearly changed as a function of CO showing an isosbestic point at 427 nm (Fig. 2A). Based on the absorbance ratio at 422 and 434 nm (*A*_422_/*A*_434_), the molar ratio of CO-hemoCD1 in the total hemoCD1 derivatives in the solution (*R*_CO_) was calculated by the following equation:

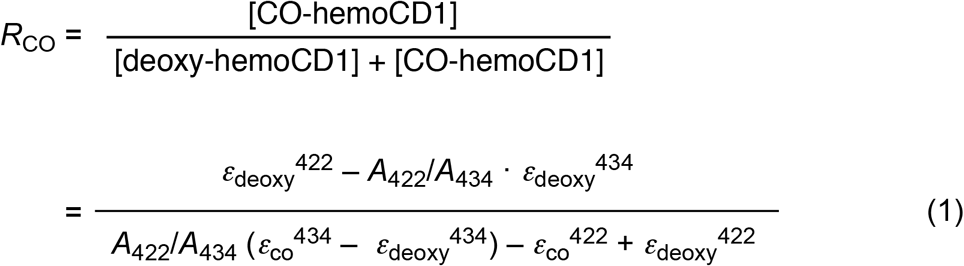

where *ε*_deoxy_^422^, *ε*_deoxy_^434^, *ε*co^422^, and *ε*_co_^434^ are the molar absorption coefficients of deoxy- and CO-hemoCD1 at 422 and 434 nm, respectively (these coefficients are listed in table S1). The derivation of eq 1 is detailed in Supplementary Materials. The total amount of hemoCD1 (*C*_total_, that is deoxy- and CO-hemoCD1 combined) was determined by the absorbance at 427 nm, the isosbestic point of deoxy- and CO-hemoCD (Fig. 2A), and its absorption coefficient (*ε*^427^, Table S1). The amount of CO (*M*_CO_) in the solution was calculated by multiplying *R*_CO_ by *C*_total_. We validated our approach by determining the *M*_CO_ values of known CO standards prepared by adding pure CO gas from a gas-tight syringe to water contained in a rubber-capped bottle without headspace. As shown in Fig. 2B, the amount of CO quantified using hemoCD1 is almost identical to the CO content in the standards (y = 0.99x, *R*^2^ = 0.999). Due to the limit of absorption detection, approximately 400 pmol was the minimum value for detection of CO. The slope of the standard curve was 1 with intercept 0; therefore, the amount of CO in solution can be directly determined using eq 1 without preparing a standard curve for each detection assay.

**Fig. 2.**
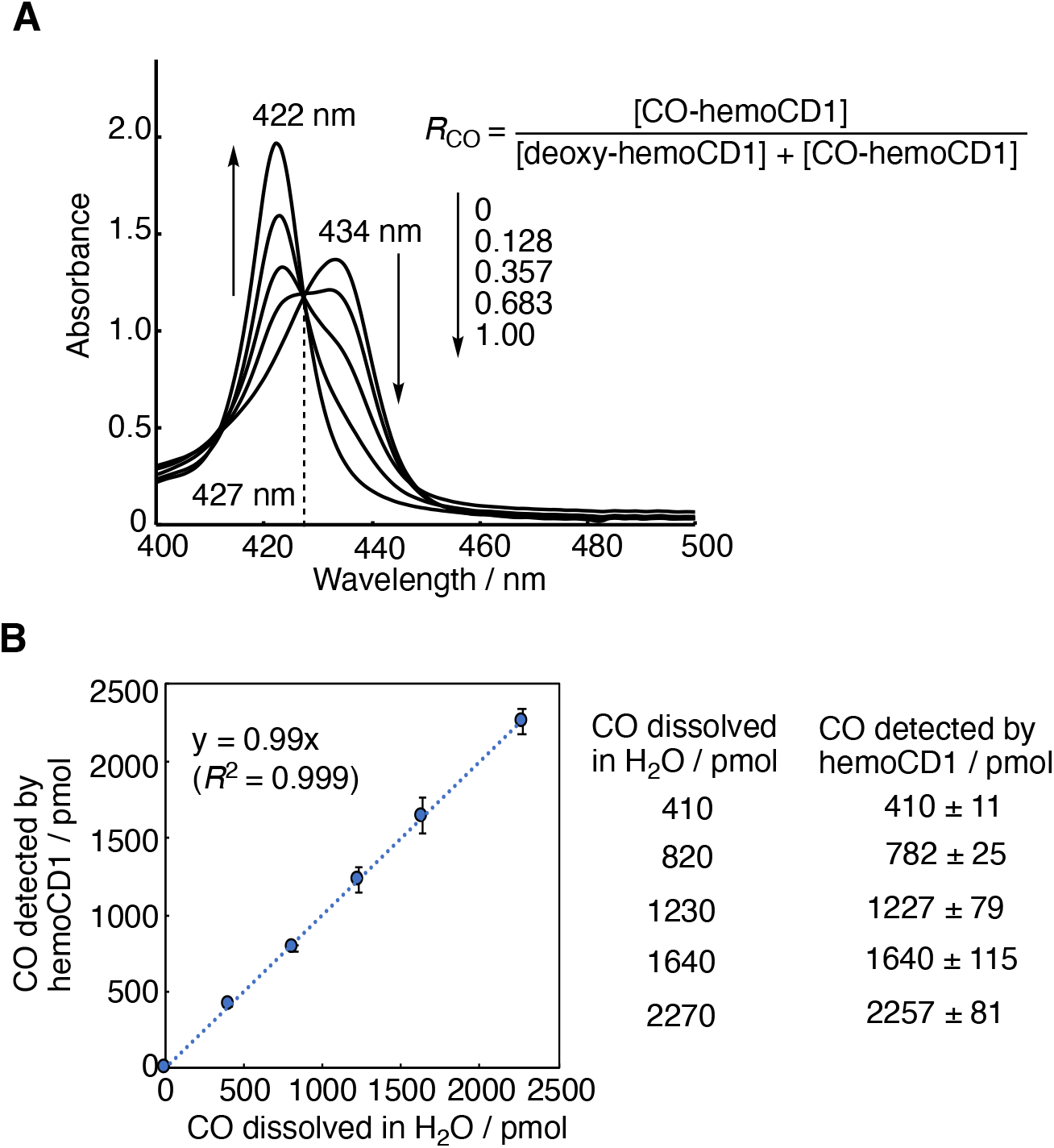
Spectroscopic quantification of CO in aqueous solution using hemoCD1. (**A**) UV-vis spectra of hemoCD1 (6.0 μM) after exposure to various amounts of CO in PBS (1 mL) at pH 7.4 and 25°C. *R*_CO_ values were calculated using eq 1. (**B**) The plot of known amounts of CO dissolved in water versus the quantified values of CO determined by the hemoCD1 assay. Data are shown as mean ± SE (*n* = 3).

### Using the new hemoCD1 assay to quantify endogenous CO levels in tissues

We next applied our new assay to determine the amount of endogenous CO contained in rat tissues. Samples (5–20 mg) collected from different organs/tissues were homogenized in PBS (0.5 mL). Then, deoxy-hemoCD1 was added to the tissue suspension followed by sonication (10 s x 2) (see Fig. 3A for methodological details). After sonication, samples were centrifuged for 15 min to obtain a clear supernatant (see picture in Fig. 3A), which was filtered through a 0.45 μm filter. Samples were then measured by UV-vis absorption spectrometry, yielding a clear Soret band as exemplified by the liver sample shown in Fig. 3B (red line). Controls were treated in the same manner without adding deoxy-hemoCD1 and their absorbances at 422 and 434 nm (Fig. 3B, blue line) were subtracted from the absorbances of samples added with deoxy-hemoCD1. From these values, *R*_CO_ was calculated using eq 1. The total content of hemoCD1 (*C*_total_) determined from the absorbance of the isosbestic point at 427 nm was consistent with the amount of deoxy-hemoCD1 initially added to the sample. This indicates that hemoCD1 was not lost during sample treatment, *i.e*., during sonication, centrifugation, and filtration. From *R*_CO_ and *C*_total_, the amount of CO contained in the tissues (*M*_CO_) was determined.

**Fig. 3.**
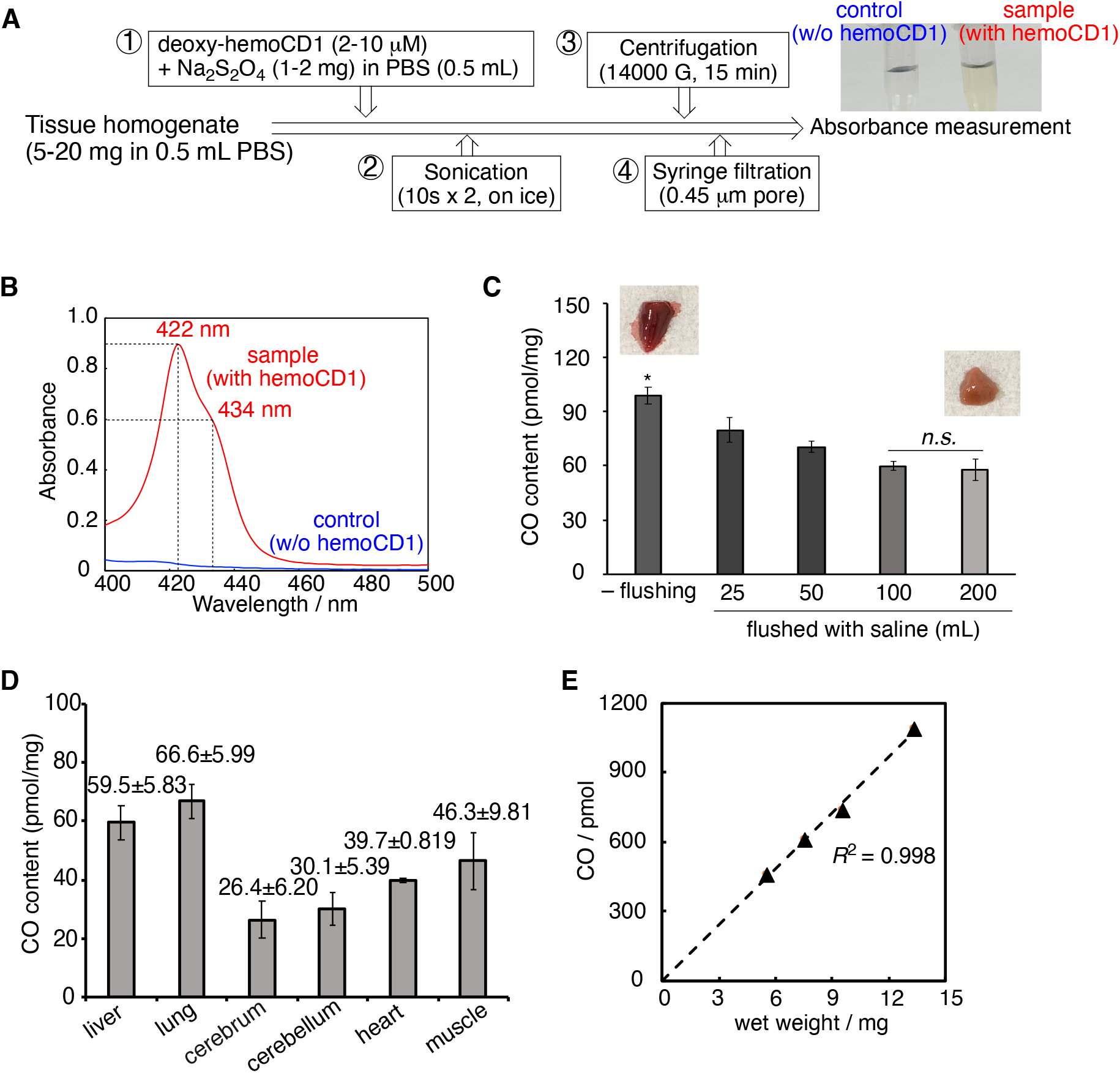
Quantification of CO in rat tissues. (**A**) Experimental procedure describing the various steps of the hemoCD1 assay for measuring CO in tissue samples collected from different rat organs. The picture shows clear supernatant solutions of tissue sonicates without (control) and with deoxy-hemoCD1. (**B**) Typical representative spectra of supernatant solutions of liver sample and control obtained at the end of the hemoCD1 assay. (**C**) Amounts of CO quantified in liver tissues without (–) or following flushing with 25–200 mL saline. Each bar represents the mean ± SE (*n* = 3–6). Statistical significance, \**p* < 0.05 versus 200 mL flushed organs; not significant, *n.s*. (**D**) Content of endogenous CO (pmol/mg, wet weight) detected in different organs using the hemoCD assay. Each bar represents the mean ± SE (*n* = 3–6). (**E**) Plot of the wet weight of liver tissue versus the amount of CO detected.

Because tissue samples contain blood, some residual CO-Hb could still be present in our samples leading to overestimation of CO levels in tissues. Therefore, we compared the CO content without and with flushing organs and tissues with saline. As shown in Fig. 3C and Fig. S9, along with a discoloring of tissues as flushing was conducted, the amount of detected CO decreased (see Fig. 3C for liver, and Fig. S9 for lung, muscle, and brain). Even though the CO content was unchanged after flushing with 100 or 200 mL saline, 200 mL saline was chosen to ensure complete flushing of organs. The varying amounts of endogenous CO (pmol/mg tissue, wet weight (ww)) measured in the different organs are reported in Fig. 3D. The amount of CO showed a linear correlation with the mass of tissues (Fig. 3E and Fig. S10), supporting the accuracy of the method quantifying CO. As for the spleen, the amount of CO could not be accurately estimated because of interference with endogenous heme-related pigments which could not be adequately removed by the treatment with Na_2_S_2_O_4_ and affected the hemoCD1 spectrum. Because hemoCD1 is stable at different pH (pH 4–10) (*28*), under room light, and in the presence of biological reactive species such as ROS, NO, H_2_S, and glutathione, the amount of CO in tissues quantified by hemoCD1 was unaffected by these external factors (Fig. S11).

We also compared our CO detection assay with a GC technique equipped with a thermal conductivity detector (TCD), following a methodology described in the literature (*16*). In GC analysis, processed tissue samples need to be oxidized by sulfosalicylic acid to liberate CO before measuring gaseous CO in the headspace (*16–19*). As shown in Fig. S12, 17.5 ± 0.6 and 12.1 ± 3.1 pmol/mg of CO were detected in liver tissue without or with flushing of the organ with 200 mL saline, respectively. The amount of CO detected in tissue by GC was consistent with data previously reported (*48*). On the other hand, we detected considerably more CO using hemoCD1 (98.7 ± 17.5 and 57.8 ± 3.0 pmol/mg). The underestimation of CO by the GC method is probably due to residual CO that is strongly bound to the tissue and can be hardly released by the denaturating agent. The assay using hemoCD1 is capable of detecting more CO in tissues than GC, even though there is still no evidence that all of CO in the tissue was detected by the assay.

### Using the hemoCD1 assay to assess the temporal distribution of CO in blood and tissues after administration of CO gas

CO present in blood and organs was measured at 5, 10, and 20 min after exposure of rats to air containing 400 ppm CO gas by inhalation (see protocol in Fig. 4A and Fig. S13). HemoCD1 was used to assess CO in tissues. Blood CO-Hb levels were measured by a blood analyzer (*49*). As expected, during the 20 min exposure to CO gas, CO-Hb in blood linearly increased to 15.3% and 12.1% in arterial and venous blood from left (LV) and right ventricles (RV), respectively (Fig. 4B). Conversely, the amounts of CO in tissues rapidly increased after 5 min exposure to CO and then reached a plateau at 10 min (Fig. 4C). Thus, it appears that, compared to Hb, the tissue has a limited capacity to store CO. We then took the same tissue samples from rats inhaled with CO gas, placed them *ex vivo* under a CO atmosphere for 1 h, and quantified the CO content. As shown in Fig. 4D, the tissues could store more CO by adding CO *ex vivo*. These results indicate that the capacity of the tissues to store CO did not reach saturation during continuous CO inhalation *in vivo*. Intriguingly, these data also suggest that the high capacity of Hb for CO storage during inhalation may confer protection of tissues against CO toxicity. This may be due, among other possibilities, to the ability of Hb to extract CO from the tissues.

**Fig. 4.**
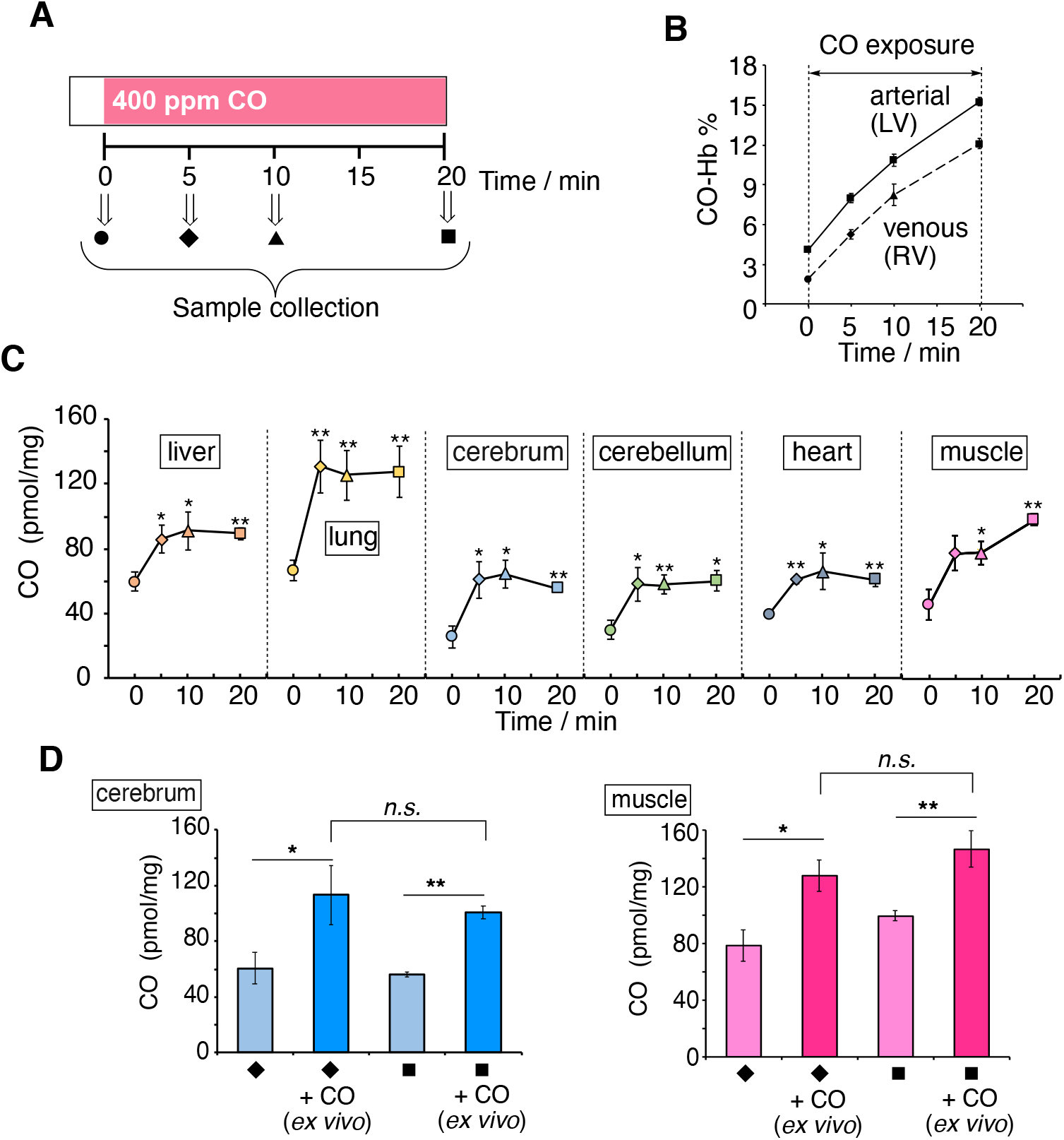
Kinetic studies of CO levels in blood and tissues after exposure to CO in rats. (**A**) Anesthetized rats were exposed to CO inhalation (400 ppm) and samples collected at different times as shown. (**B**) Changes in CO-Hb (%) in arterial and venous blood as a function of time. (**C**) Tissue CO contents as a function of time. Each plot represents the mean ± SE (*n* = 3–6). Statistical significance, \**p* < 0.05, \*\**p* < 0.01 versus *t* = 0. (**D**) Amount of CO in muscle and cerebrum before and after purging CO gas *ex vivo*. Samples collected at 5 or 20 min were placed under CO atmosphere for 1 h and then assayed for CO content. Each bar represents the mean ± SE (*n* = 3). Statistical significance, \**p* < 0.05, \*\**p* < 0.01; not significant, *n.s*.

To demonstrate this hypothesis, we measured CO accumulated in hepatocytes incubated for 2 h with a CO-releasing molecule (CORM-401E), followed by replacement of medium in the presence or absence of Hb (Fig. S14). Notably, hepatocytes exposed to CORM-401E exhibited a significant increase in intracellular CO, whereas CO levels were equivalent to control values after treatment with CORM401-E followed by oxy-Hb, indicating that CO initially accumulated in cells was subsequently transferred to Hb. These results suggest that Hb is capable of removing CO stored in cells, which was confirmed by the CO quantification assay using hemoCD1, and that this mechanism could also occur *in vivo*, supporting a possible protective role of Hb against CO intoxication (see Discussion for detail).

Next, the kinetic profiles of CO levels in blood and tissues were studied in rats first exposed to 400 ppm CO for 5 min followed by ventilation with air or pure O_2_ for 60 min (Fig. 5A). As expected, CO-Hb levels in arterial (LV) and venous blood (RV) gradually decreased after the two treatments, with O_2_ ventilation being more effective than air (Fig. 5B). However, a significant amount of CO still remained in several tissues, especially in the brain (cerebrum and cerebellum), even after ventilation with O_2_ (Fig. 5C). Thus, once CO is accumulated in tissues it is not easily eliminated by standard treatments used against CO intoxication.

**Fig. 5.**
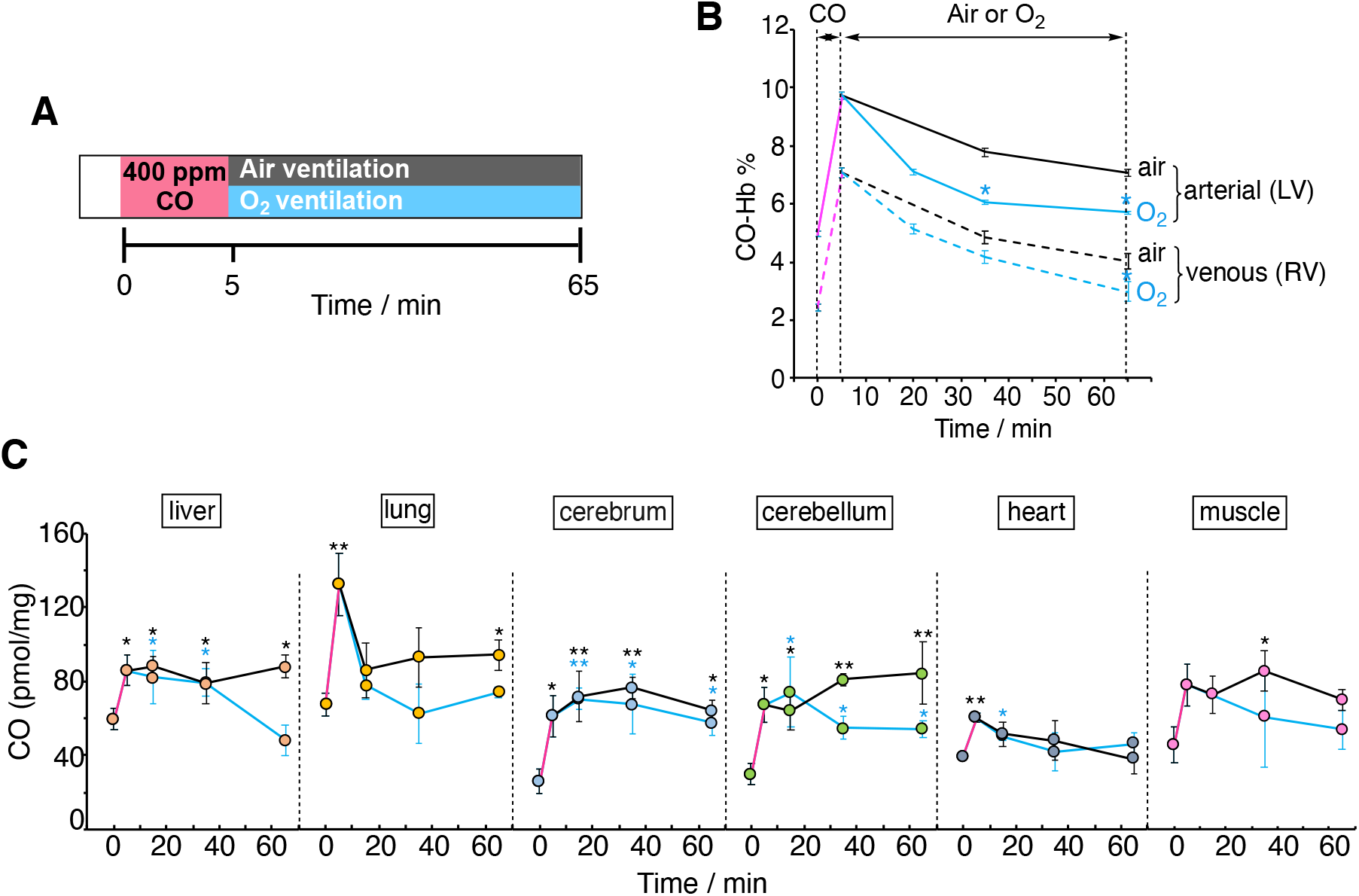
Effect of normobaric air/O_2_ ventilation on CO levels in blood and tissues after exposure to CO in rats. (**A**) Anesthetized rats were exposed to CO inhalation (400 ppm) for 5 min followed by either air (black) or O_2_ ventilation (blue) as indicated. (**B**) Changes in CO-Hb (%) in arterial and venous blood as a function of time. Each plot represents mean ± SE (*n* = 5–14). Statistical significance, \**p* < 0.05, versus air ventilation. (**C**) Amounts of CO measured under the experimental conditions described in (**A**). The plots connected by black and blue lines represent the data obtained under air and O_2_ ventilation, respectively. Each bar represents mean ± SE (*n* = 3–6). Statistical significance, \**p* < 0.05, \*\**p* < 0.01; not significant, *n.s*. versus *t* = 0.

### Testing hemoCD1 as an antidote against CO intoxication

Finally, we tested whether oxy-hemoCD1 could act as an effective CO-removal agent *in vivo* during CO intoxication (Fig. 6). At first, three types of experimental protocols were used as shown in Fig. 6A. In protocols ***I*** and ***II***, rats were subjected to 5 min CO inhalation followed by an intravenous (i.v.) injection of oxy-hemoCD1 during air (***I***) or O_2_ ventilation (***II***). In the third protocol (***III***), CO inhalation was followed by O_2_ ventilation and, 30 min later, by injection of oxy-hemoCD1. Immediately after the injection, oxy-hemoCD1 was started to be excreted in the urine in the form of CO-hemoCD1 (Fig. S15) and the pharmacokinetic study on oxy-hemoCD1 has been reported elsewhere (*27,36*). The data in Fig. 6A show that oxy-hemoCD1 infusion during air ventilation (***I***) fastens the return of CO-Hb to basal levels, while during O_2_ ventilation (***II***) the effect of hemoCD1 infusion is less pronounced. On the other hand, the infusion of oxy-hemoCD1 elicited a marked reduction in CO accumulation in the brain (cerebrum and cerebellum, see Fig. 6A) and other tissues (Fig. S16). The combination of oxy-hemoCD1 with O_2_ ventilation (***II*** and ***III***) was especially effective in removing CO from the brain.

**Fig. 6.**
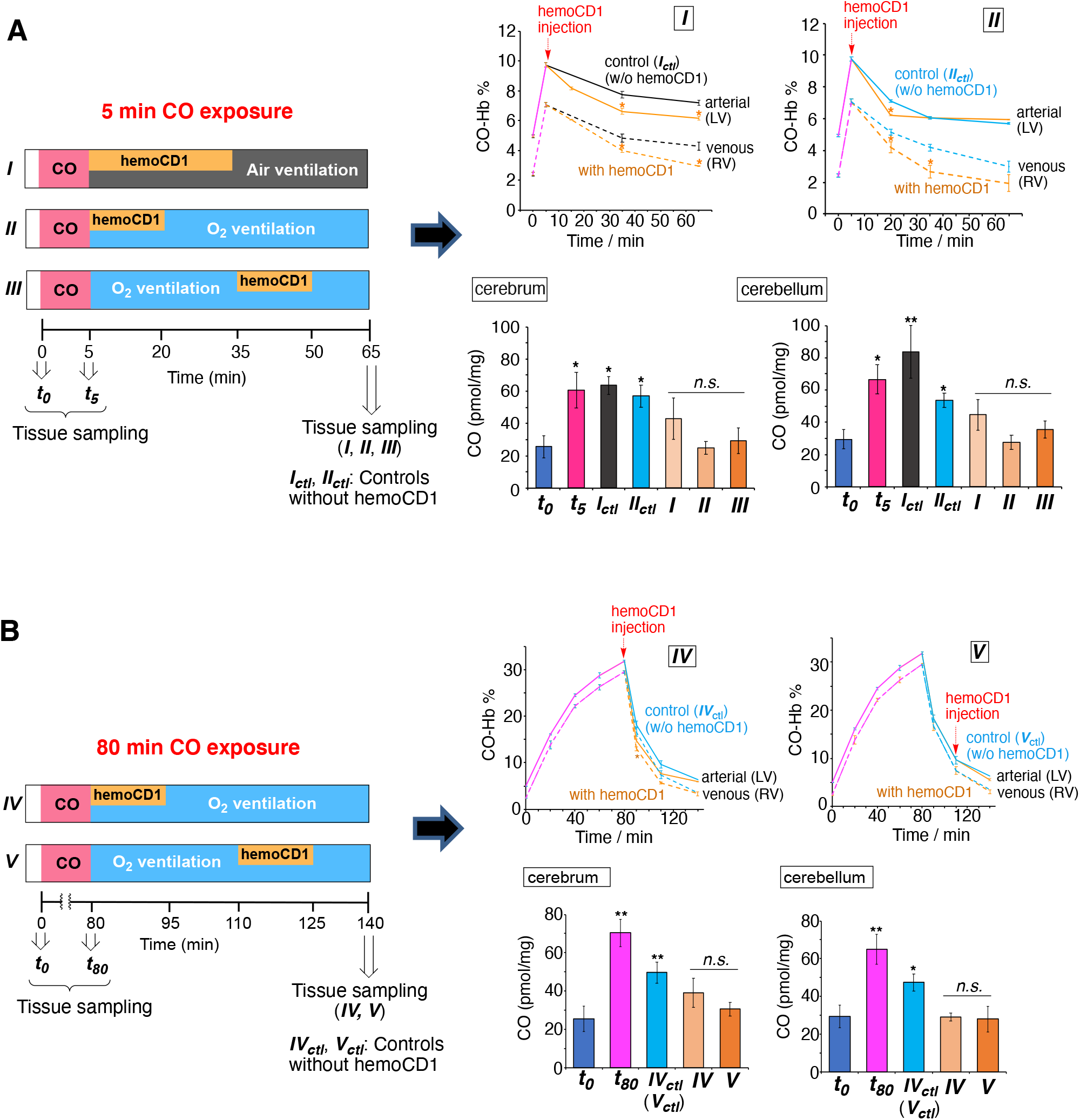
Effect of normobaric air/O_2_ ventilation in combination with hemoCD1 injection on CO levels in blood and tissues after exposure to exogenous CO in rats. (**A**) Anesthetized rats were exposed to CO inhalation (400 ppm) for 5 min followed by either air or pure O_2_ ventilation in combination with intravenous hemoCD1 infusion (1.4 ± 0.2 mM, 2.5 mL in PBS) as indicated by the three different protocols: ***I***, oxy-hemoCD1 was infused for 30 min under room air ventilation; ***II***: oxy-hemoCD1 was infused for 15 min under pure O_2_ ventilation; ***III***: O_2_ ventilation was conducted for 30 min before infusion of oxy-hemoCD1 for 15 min. The right panels show the changes in CO-Hb (%) in arterial (LV) and venous (RV) blood and the CO levels detected in the cerebrum and cerebellum samples collected as indicated. (**B**) Anesthetized rats were exposed to CO inhalation (400 ppm) for 80 min followed by O_2_ ventilation in combination with intravenous hemoCD1 infusion (3.0 ± 0.2 mM, 2.5 mL in PBS) as indicated by the two different protocols: ***IV***, oxy-hemoCD1 was infused for 15 min under pure O_2_ ventilation; ***V***: O_2_ ventilation was conducted for 30 min before infusion of oxy-hemoCD1 for 15 min. The right panels show the changes in CO-Hb (%) in LV and RV blood and the CO levels detected in the cerebrum and cerebellum samples collected as indicated. Each plot for CO-Hb (%) represents mean ± SE (*n* = 3–14). Each bar for CO (pmol/mg) represents the mean ± SE (*n* = 3–6). Statistical significance, \**p* < 0.05, \*\**p* < 0.01; not significant, *n.s*. versus ***t*_0_**.

To simulate more closely a state of severe CO intoxication, the effect of oxy-hemoCD1 administration was further tested in rats after inhalation of 400 ppm CO for 80 min. Under these conditions, CO-Hb levels reached 30% (Fig. 6B). Unexpectedly, the amount of CO accumulated in tissues was not significantly increased compared to that measured after shorter CO inhalation times (5 and 20 min) (Fig. 6A, 6B, and Fig. S17). This observation is consistent with the scenario that a high capacity for CO storage of circulating Hb impedes accumulation of excess CO in tissues (see Discussion for detail). Also in this case, the significant amount of residual CO detected in the brain after O_2_ ventilation for 60 min (***IV_ctl_***/***V_ctl_***) was effectively reduced by i.v. injection with oxy-hemoCD1 (***IV*** and ***V***).

## Discussion

In this study, we report on the ability of hemoCD1 to effectively remove CO from tissues following CO exposure in rats. Our data show that inhalation of CO gas rapidly increases CO-Hb and CO accumulation in tissues *in vivo*. While air or O_2_ ventilation is capable of restoring CO-Hb to basal levels, elimination of CO from tissues is hard to achieve, especially in the brain. However, infusion of hemoCD1 during the ventilation treatments efficiently eliminates CO from tissues, uncovering a very useful property of hemoCD1 as a CO removing agent to combat CO intoxication. We further reveal how tissues rapidly reach a plateau in CO content during CO exposure while CO-Hb levels in blood continue to rise over time. This phenomenon raises important questions on the role of Hb as a potential molecular shield that defends tissues against accumulation of dangerous levels of CO. The data on CO content in tissues and organs were generated using a new and simplified CO detection method based on hemoCD1, which was thoroughly investigated and validated in the present study.

Our data show that the CO-hemoCD1 complex, unlike CO-Hb, is extremely stable and hardly decomposes under extreme conditions such as high pressures of N2 and O_2_, or in the presence of oxidants (H_2_O_2_). The facts that hemoCD1: 1) is not cell-permeable (*39*) and not easily denatured by other chemicals; 2) can be isolated from other biocomponents; and that 3) its O_2_ binding affinity is moderate (Table 1) enabling CO to easily replace O_2_, make it a unique molecule with suitable properties as a CO scavenging agent in the biological environment. Another important property is that hemoCD1 also exhibits a much higher CO binding affinity compared to NO and H_2_S.

**Table 1.**
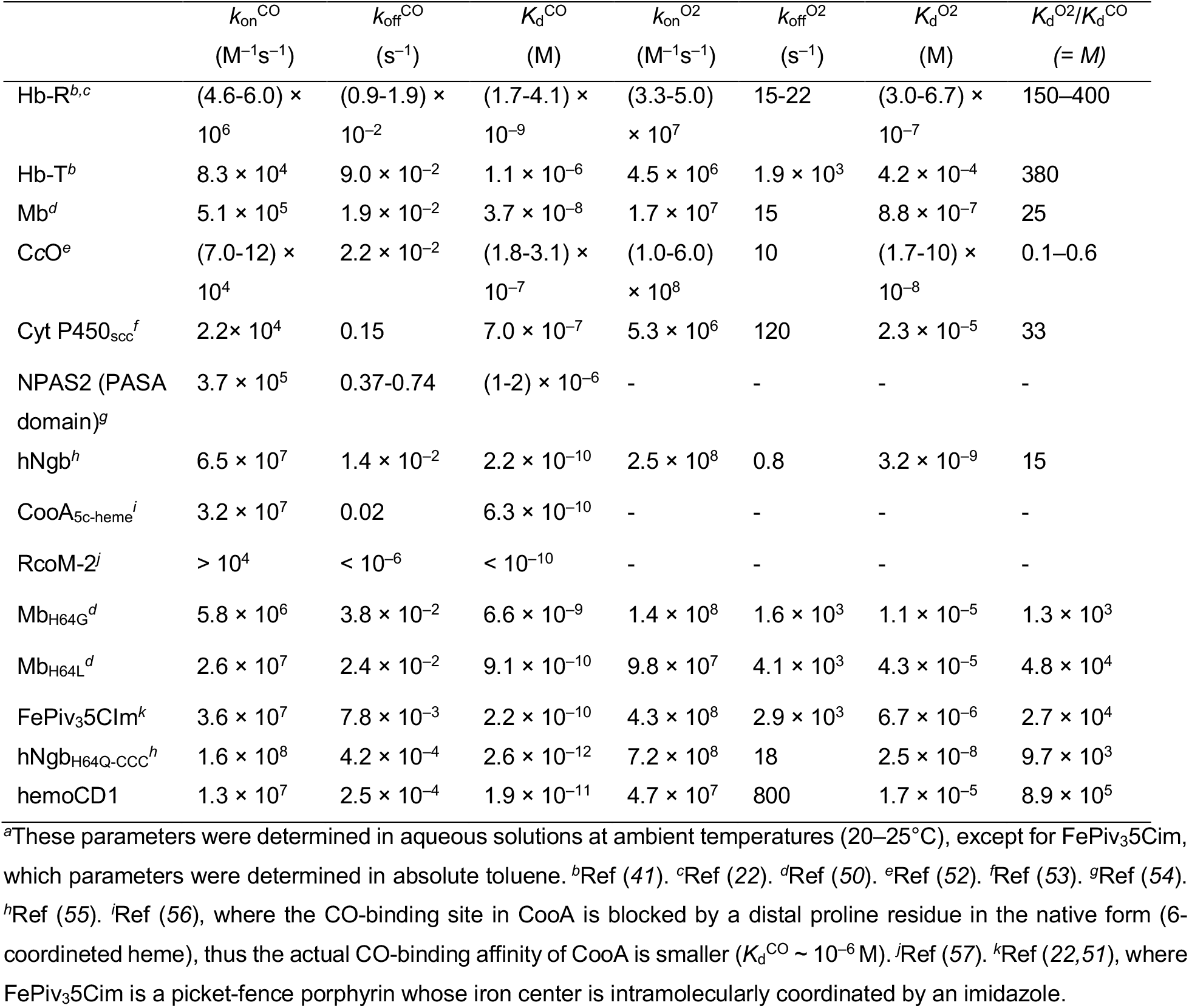
Kinetic and thermodynamic parameters for O_2_ and CO bindings of hemoCD1 and related hemoproteins and model compounds.^*a*^

Notably, hemoCD1 binds CO with much stronger affinity than Hb, Mb, C*c*O, and other hemoproteins found in nature (Table 1). The CO binding affinity tends to be higher in CO-sensing proteins such as neuroglobin (Ngb), NPAS2, CooA, and RcoM; in the case of hemoCD1, the CO affinity is still one order of magnitude higher than that of CO-sensing proteins. The kinetic parameters indicate that the high CO binding affinity of hemoCD1 is ascribed to the slow CO off rate (*k*off^CO^ = 2.5 × 10^−4^ s^−1^). The hydrophobic environment provided by per-*O*-methyl-β-CD in CO-hemoCD1 tightly holds a hydrophobic CO molecule on the iron(II) center of hemoCD1 as discussed elsewhere (*28*). In native Hb and Mb, O_2_/CO binding sites are surrounded by a steric and polar amino acid residue, called distal His, which reduce the CO binding affinities relative to O_2_ (*22,41*). Distal mutants such as H64G and H64L Mb showed much higher CO affinity than that in native Mb, resulting in large *M* values (see Table 1) (*50*). In synthetic iron(II)porphyrins such as hemoCD1 and FePiv_3_5Cim (*51*), which have no distal functional groups, the CO binding affinity is much higher than that of Mb and Hb (*22*), although the lipophilic model cannot be used as CO-scavengers in biological media.

In the present study, we refined and optimized an assay using hemoCD1 for the specific detection of CO in tissues and organs. We previously measured CO in cells using oxy-hemoCD1 (*39*), which can detect CO via ligand exchange reaction from O_2_ to CO but necessitated three troublesome processes: 1) a gel filtration step to remove excess reductant (Na2S2O4) for the preparation of oxy-hemoCD1; 2) an ultrafiltration process to separate hemoCD1 from biological contaminants; and 3) a CO gas bubbling step to convert all the hemoCD1 forms to CO-hemoCD1 for determining the final concentration of hemoCD1 (*C*_total_) in the sample. Here we have improved our method by eliminating the gel filtration step since the amount of CO in tissues is determined based on the absorbance ratio at 422 and 434 nm in the presence of excess dithionite. Dithionite does not affect the CO scavenging properties of hemoCD1 and facilitates the precipitation of bio-components present in tissue samples, allowing to obtain clear solutions for absorbance measurements after centrifugation. The ultrafiltration process was thus also omitted. Furthermore, the ratiometric assay using *A*_422_/*A*_434_ and *A*_427_ enables to determine *M*_CO_ in a single spectrum measurement, eliminating the final CO bubbling step. We also confirmed that the new assay can be used for cultured cells *in vitro*, measuring the same amount of endogenous CO as previously reported (*38*). Therefore, the new assay using hemoCD1 is accurate and represents a convenient method to determine the amount of CO stored in biological samples such as cells/tissues/organs. We note that different methods to detect CO in tissues and cells have been developed by several groups (*10,16–19,58–66*). CO detection in cells has been first achieved with the fluorescent probe COP-1 (*58*) and subsequently other Pd-based CO sensitive probes were synthesized (*60–63*) but none of them can quantify CO in samples. Although laser spectroscopy and radio-isotope methods have been reported (*64–66*), GC remains the most accessible and common technique to quantify CO. As we demonstrated in this study, CO quantification in tissues was underestimated by GC analysis compared to our method, highlighting once more the sensitivity of the hemoCD1 assay.

Our study on the kinetic profiles of CO in tissues and blood reveals interesting dynamics of CO distribution. First, the amount of CO stored in tissues quickly reached a plateau during CO inhalation but the *ex vivo* experiments revealed that tissue samples can store more CO. Secondly, despite the plateau observed in tissues, CO-Hb in blood linearly increased during CO exposure. What kind of mechanism could explain this dynamic? We postulate a scenario according to which CO reaching the tissues would gradually transfer to Hb in RBC where it would be stored as CO-Hb before elimination through the lungs. The schematic representation of uptake and elimination of CO is proposed in Fig. 7. In normal conditions, endogenous CO is continuously produced in every cell, some of which is stored in tissues and gradually eliminated through RBC (Fig. 7A). Following inhalation, CO distributes in the body: 95% readily binds to Hb in blood (*11*) and the rest that is not captured by RBC diffuses to tissues (Fig. 7B). As the CO binding affinity of Hb is much higher than those of Mb and C*c*O (Table 1), which are the major CO storage components in tissues, it is reasonable to assume that CO flows from tissues to Hb in RBC. Thus, continuous CO inhalation leads to a further increase in CO-Hb, while CO in tissues reaches a certain plateau, *i.e*. an equilibrium state (Fig. 7C). This is corroborated by our two CO exposure protocols, since the amount of CO accumulated in tissues was very similar but CO-Hb reached levels of 10 and 30% after inhalation of CO for 5 and 80 min, respectively. The CO flow mechanism suggests that Hb in RBC plays a role in protecting surrounding tissues from CO toxicity. In fact, as demonstrated in the dog study (*14*) mentioned in the Introduction, high CO-Hb itself is not toxic to the animals and several studies highlight that it is the CO diffused to the tissues and not the fraction bound to Hb that is the principal cause of CO poisoning (*8,14,55*). Our data showing that exogenous Hb can capture CO previously accumulated in hepatocytes *in vitro* support this proposed mechanism. Likewise, the fact that CO accumulation in organs *ex vivo* is higher than that measured after CO exposure *in vivo* indicates that, in the absence of Hb, more CO can reach the tissue. Our results also show that O_2_ ventilations after CO inhalation is more efficient than air in removing CO from blood. However, significant amounts of CO still remained in several tissues even after 60 min O_2_ ventilation. In particular, the residual CO in the brain might be the cause of CO-associated delayed neurological sequelae (DNS) (*8,67*) as up to 40% of subjects surviving acute CO poisoning may develop DNS in 2 to 40 days. This may lead to memory loss, movement disorders, and Parkinson-like syndrome among others (*67*). The pathophysiology of CO poisoning and subsequent DNS is poorly understood, although CO-induced mitochondrial dysfunction is likely a cause of brain injury (*8,68–70*). Our data on CO quantification indicates that CO, once stored in the brain, is relatively hard to eliminate and normobaric O_2_ ventilation is insufficient to completely remove CO from this tissue.

**Fig. 7.**
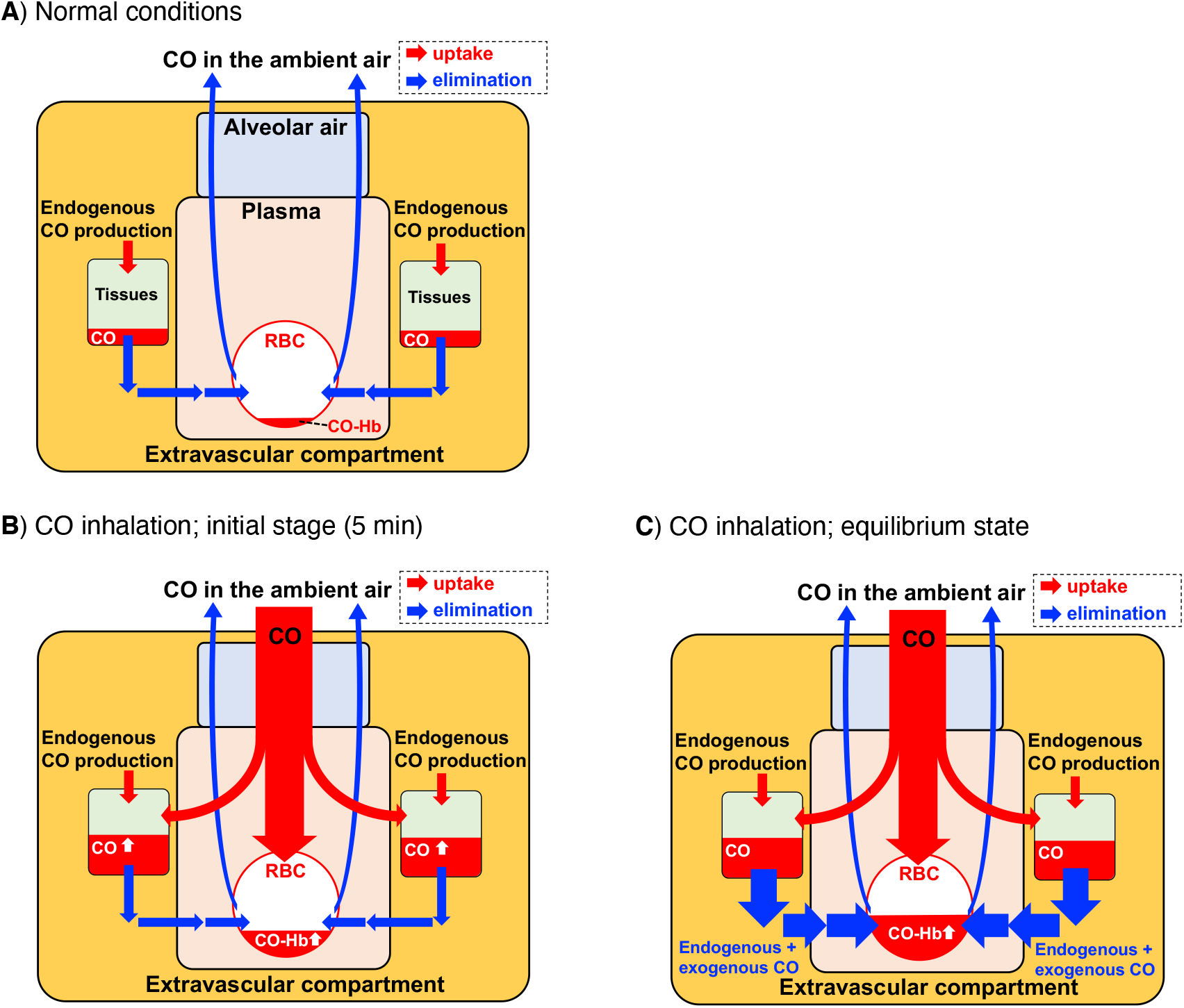
Proposed mechanism of CO compartmentalization under normal conditions and during CO inhalation. (**A**) Normal conditions. Endogenous CO continuously produced in cells is stored in tissues, diffuses to Hb, and is exhaled. (**B**) Initial stage of CO inhalation. Inhaled CO forms CO-Hb in RBC and diffused to tissues. (**C**) Equilibrium state during CO inhalation. CO accumulated in tissues gradually transfers to Hb in RBC based on the higher CO affinity of Hb versus intracellular CO targets (see Table 1 and text for details). The compartment models are based on our data and Refs (*9*) and (*17*).

Thus, in the last part of our study we tested the potential use of hemoCD1 as an antidote for CO intoxication considering that O_2_ ventilation is the only method practically used as a therapy. Recently, a neuroglobin mutant (Ngb-H64Q-CCC) (*52,55*), which showed much higher CO binding affinity than Hb and hemoCD1 (Table 1), has been proposed as an injectable type of CO antidote. Our data demonstrate that i.v. injection of oxy-hemoCD1 was effective in decreasing CO levels in the brain of CO-treated rats. Since our previous study (*38*) showed no evidence that hemoCD1 diffuses through the blood brain barrier, we speculate that an equilibrium diffusion of CO to hemoCD1 in plasma effectively reduces CO in the brain. In any case, oxy-hemoCD1 injection reduced CO-Hb in blood (*36*), which may boost the elimination of CO in tissues by circulating Hb.

In conclusion, we propose the use of hemoCD1 as a useful CO scavenger for quantifying and removing exogenous CO in tissues. Our study emphasizes: (1) the high affinity of hemoCD1 for CO and the chemical stability of the CO-hemoCD1 complex, demonstrating the suitability of hemoCD1 as a sensitive CO scavenger; (2) that the ratiometric absorbance assay using hemoCD1 in the presence of dithionite is a very simple and sensitive method for quantifying CO contained in tissues; (3) an unrecognized role of Hb as a predominant CO storage system *in vivo*, preventing accumulation of excess CO in tissues; (4) that injection of oxy-hemoCD1 to CO-exposed rats acts as adjuvant to air/O_2_ ventilation to effectively and rapidly remove CO accumulated in tissues, including the brain. We believe that the data presented herein will stimulate further discussions on the mechanisms underlying CO poisoning.

## Materials and Methods

Fe^II^TPPS and Py3CD are synthesized in our laboratory according to the literature method (*27,28*). All animal experiments were approved by the Institutional Review Board of Tokai University and the Guidelines for Animal Experiments of Doshisha University. An extended materials and methods section is available in Supplementary Information. This includes details on sample preparation, animal experiments, and the assay for quantification of CO using hemoCD1.

## Acknowledgments

This work was financially supported by MEXT/JSPS KAKENHI (Grant No. JP15H02569, JP17H02208, JP18KK0156, JP19K22260, JP19K22972, JP20H02871), the MEXT-Supported Program for the Strategic Research Foundation at Private Universities (2015–2019), the Takeda Science Foundation, the NOVARTIS Foundation (Japan) for the Promotion of Science, the Suntory Foundation for Life Sciences, and the JGC-S Scholarship Foundation. Q.M. deeply thanks the support of Otsuka Toshimi Scholarship Foundation and a Visiting Fellowship from IMRB/Inserm. RF and RM are supported by a grant from the Agence National de la Recherche (ANR-19-CE18-003201SWEET-CO).

## Author Contributions

Q.M., A.T.K., S.M., and H.K. performed experiments and analyzed data; Q.M., R.M., R.F., and H.K. conceived the concept of the study and wrote the manuscript. The authors declare no competing interest.

## References and Notes

1. T. M. Dydek, “Investigating carbon monoxide poisonings.” in Carbon Monoxide Poisoning, D. G. Penney, Ed. (CRC-Taylor and Francis Press, NY, 2007), Chapter 12.

2. R. Motterlini, R. Foresti, Biological signaling by carbon monoxide and carbon monoxide-releasing molecules. Am. J. Physiol. Cell Physiol. 312, C302–C313 (2017).

3. T. Shimizu, D. Huang, F. Yan, M. Stranava, M. Bartosova, V. Fojtíková, M. Martínková, Gaseous O2, NO, and CO in signal transduction: Structure and function relationships of heme-based gas sensors and heme-redox sensors. Chem. Rev. 115, 6491–6533 (2015).

4. C. Szabo, Gasotransmitters in cancer: from pathphysiology to experimental therapy. Nat. Rev. Drug Discovery 15, 185–203 (2016).

5. J. M. Fukuto, S. J. Carrington, D. J. Tantillo, J. G. Harrison, L. J. Ignarro, B. A. Freeman, A. Chen, D. A. Wink, Small molecule signaling agents: The integrated chemistry and biochemistry of nitrogen oxides, oxides of carbon, dioxygen, hydrogen sulfide, and their derived species. Chem. Res. Toxicol. 25, 769–793 (2012).

6. G. B. Ray, X.-Y. Li, J. A. Ibers, J. L. Sessler, T. G. Spiro, How far can proteins bend the FeCO unit? Distal polar and steric effects in heme proteins and models. J. Am. Chem. Soc. 116, 162–176 (1994).

7. M. Matsu-ura, F. Tani, Y. Naruta, Formation and characterization of carbon monoxide adducts of iron “twin coronet” porphyrins. extremely low CO affinity and a strong negative polar effect on bound CO. J. Am. Chem. Soc. 124, 1941–1950 (2002).

8. J. J. Rose, L. Wang, Q. Xu, C. F. McTiernan, S. Shiva, J. Tejero, M. T. Gladwin, Carbon monoxide poisoning: pathogenesis, management, and future directions of therapy. Am. J. Respir. Crit. Care Med. 195, 596–606 (2017).

9. US-EPA, “Pharmacokinetics and Mechanisms of Action of Carbon Monoxide.” in Air Quality Criteria for Carbon Monoxide, Chapter 5, pp. 5–1–5-30 (2000).

10. B. Widdop, Analysis of carbon monoxide. Ann. Clin. Biochem. 39, 378–391 (2002).

11. P. G. Flachsbart, “Exposure to ambient and microenvironmental concentrations of carbon monoxide” in Carbon Monoxide Poisoning, D. G. Penney, Ed. (CRC-Taylor and Francis Press, NY, 2007), Chapter 2.

12. L. K. Weaver, R. O. Hopkins, K. J. Chan, S. Churchill, C. G. Elliott, T. P. Clemmer, J. F. Orme Jr, F. O. Thomas, A. H. Morris, Hyperbaric Oxygen for Acute Carbon Monoxide Poisoning. N. Engl. J. Med. 347, 1057–1067 (2002).

13. C. Tomaszewski, “The case for the use of hyperbaric oxygen therapy in carbon monoxide poisoning” in Carbon Monoxide Poisoning, D. G. Penney, Ed. (CRC-Taylor and Francis Press, NY, 2007), Chapter 2.

14. L. R. Goldbaum, R. G. Ramirez, K. B. Absalon, What is the mechanism of carbon monoxide toxicity? Aviat. Space Environ. Med. 46, 1289–1291 (1975).

15. N. B. Hampson, N. M. Hauff, Risk factors for short-term mortality from carbon monoxide poisoning treated with hyperbaric oxygen. Crit. Care Med. 36, 2523–2527 (2008).

16. H. J. Vreman, R. J. Wong, T. Kadotani, T. K. Stevenson, Determination of carbon monoxide (CO) in rodent tissue: Effect of heme administration and environmental CO exposure. Anal. Biochem. 341, 280–289 (2005).

17. R. F. Coburn, Endogenous Carbon monoxide production and body CO stores. Acta Med. Scand. Suppl. 472, 269–282 (1967).

18. H. J. Vreman, R. J. Wong, D. K. Stevenson, J. E. Smialek, D. R. Fowler, L. Li, R. D. Vigorito, H. R. Zielke, Concentration of carbon monoxide (CO) in postmortem human tissues: effect of environmental CO exposure. J. Forensic. Sci. 51, 1182–1190 (2006).

19. F. J. Cronje, M. S. Carraway, J. J. Freiberger, H. B. Suliman, C. A. Piantadosi, Carbon monoxide actuates O_2_-limited heme degradation in the rat brain. Free Radic. Biol. Med. 37, 1802–1812 (2004).

20. J. P. Collman, R. R. Gagne, T. R. Halbert, J. C. Marchon, C. A. Reed, Reversible oxygen adduct formation in ferrous complexes derived from a picket fence porphyrin. Model for oxymyoglobin. J. Am. Chem. Soc. 95, 7868–7870 (1973).

21. M. Momenteau, C. A. Reed, Synthetic heme-dioxygen complexes, *Chem*. Rev. 94, 659–698 (1994).

22. J. P. Collman, R. Boulatov, C. J. Sunderland, L. Fu, Functional analogues of cytochrome *c* oxidase, myoglobin, and hemoglobin. Chem. Rev. 104, 561–588 (2004).

23. K. Shikama, The molecular mechanism of autoxidation for myoglobin and hemoglobin: A venerable puzzle. Chem. Rev. 98, 1357–1374 (1998).

24. K. Shikama, Stability properties of dioxygen-iron(II) porphyrins: an overview from simple complexes to myoglobin. Coord. Chem. Rev. 83, 73–91 (1988).

25. D.-L. Jiang, T. Aida, A dendritic iron porphyrin as a novel haemoprotein mimic: effects of the dendrimer cage on dioxygen-binding activity. Chem. Commun. 1523–1524 (1996).

26. T. Komatsu, M. Moritake, A. Nakagawa, E. Tsuchida, Self-organized lipid-porphyrin bilayer membranes in vesicular form: nanostructure, photophysical properties, and dioxygen coordination. Chem. Eur. J. 8, 5469–5480 (2002).

27. K. Kano, H. Kitagishi, M. Kodera, S. Hirota, Dioxygen binding to a simple myoglobin model in aqueous solution. Angew. Chem. Int. Ed. 44, 435–438 (2005).

28. K. Kano, H. Kitagishi, C. Dagallier, M. Kodera, T. Matsuo, T. Hayashi, Y. Hisaeda, S. Hirota, Iron porphyrin-cyclodextrin supramolecular complex as a functional model of myoglobin in aqueous solution. Inorg. Chem. 45, 4448–4460 (2006).

29. K. Kano, Y. Itoh, H. Kitagishi, T. Hayashi, S. Hirota, A supramolecular receptor of diatomic molecules (O_2_, CO, NO) in aqueous solution. J. Am. Chem. Soc. 130, 8006–8015 (2008).

30. H. Kitagishi, M. Tamaki, T. Ueda, S. Hirota, T. Ohta, Y. Naruta, K. Kano, Oxoferryl porphyrin/hydrogen peroxide system whose behavior is equivalent to hydroperoxoferric porphyrin. J. Am. Chem. Soc. 132, 16730–16732 (2010).

31. K. Kano, S. Chimoto, M. Tamaki, Y. Itoh, H. Kitagishi, Supramolecular dioxygen receptors composed of an anionic water-soluble porphinatoiron(II) and cyclodextrin dimers. Dalton Trans. 41, 453–461 (2012).

32. K. Watanabe, H. Kitagishi, K. Kano, Supramolecular iron porphyrin/cyclodextrin dimer complex that mimics the functions of hemoglobin and methemoglobin. Angew. Chem. Int. Ed. 52, 6894–6897 (2013).

33. H. Kitagishi, S. Kurosawa, K. Kano, Intramolecular oxidative *O*-demethylation of an oxoferryl porphyrin complexed with a per-*O*-methylated β-cyclodextrin Dimer. Chem. Asian J. 11, 3213–3219 (2016).

34. K. Kano, T. Ochi, S. Okunaka, Y. Ota, K. Karasugi, T. Ueda, H. Kitagishi, Preparation and function of poly(acrylic acid)s modified by supramolecular complex composed of porphinatoiron and a cyclodextrin dimer that bind diatomic molecules (O_2_ and CO) in aqueous solution. Chem. Asian. J. 6, 2946–2955 (2011).

35. H. Kitagishi, S. Negi, A. Kiriyama, A. Honbo, Y. Sugiura, A. T. Kawaguchi, K. Kano, A diatomic molecule receptor that removes CO in a living organism. Angew. Chem. Int. Ed. 49, 1312–1315 (2010).

36. H. Kitagishi, S. Minegishi, A. Yumura, S. Negi, S. Taketani, Y. Amagase, Y. Mizukawa, T. Urushidani, Y. Sugiura, K. Kano, Feedback response to selective depletion of endogenous carbon monoxide in the blood. J. Am. Chem. Soc. 138, 5417–5425 (2016).

37. H. Kitagishi, S. Minegishi, Iron(II)porphyrin–cyclodextrin supramolecular complex as a carbon monoxide-depleting agent in living organisms. Chem. Pharm. Bull. 65, 336–340 (2017).

38. S. Minegishi, I. Sagami, S. Negi, K. Kano, H. Kitagishi, Circadian clock disruption by selective removal of endogenous carbon monoxide. Sci. Rep. 8, 11996 (2018).

39. S. Minegishi, A. Yumura, H. Miyoshi, S. Negi, S. Taketani, R. Motterlini, R. Foresti, K. Kano, H. Kitagishi, Detection and removal of endogenous carbon monoxide by selective and cell-permeable hemoprotein-model complexes. J. Am. Chem. Soc., 139, 5984–5991 (2017).

40. E. Wilhelm, R. Battino, R. J. Wilcock, Low-pressure solubility of gases in liquid water, *Chem*. Rev. 77, 219–262 (1977).

41. C. E. Cooper, Nitric oxide and iron proteins. Biochim. Biophys. Acta 1411, 290–309 (1999).

42. K. Watanabe, T. Suzuki, H. Kitagishi, K. Kano, Reaction between a haemoglobin model compound and hydrosulphide in aqueous solution. Chem. Commun. 51, 4059–4061 (2015).

43. K. Karasugi, H. Kitagishi, K. Kano, Modification of a dioxygen carrier, hemoCD, with PEGylated dendrons for extension of circulation time in the bloodstream. Bioconjugate Chem. 23, 2365–2376 (2012).

44. S. M. Mitrione, P. Villalon, J. D. Lutton, R. D. Levere, N. G. Abraham, Inhibition of Human Adult and Fetal Heme Oxygenase by New Synthetic Heme Analogues. Am. J. Med. Sci. 296, 180–186 (1988).

45. C. Shirataki, O. Shoji, M. Terada, S.-i. Ozaki, H. Sugimoto, Y. Shiro, Y. Watanabe, Inhibition of Heme Uptake in Pseudomonas aeruginosa by its Hemophore (HasAp) Bound to Synthetic Metal Complexes. Angew. Chem. Int. Ed. 53, 2862–2866 (2014).

46. K. Watanabe, H. Kitagishi, K. Kano, Supramolecular ferric porphyrins as cyanide receptors in aqueous solution. ACS Med. Chem. Lett. 2, 943–947 (2011).

47. T. Yamagiwa, A. T. Kawaguchi, T. Saito, S. Inoue, S. Morita, K. Watanabe, H. Kitagishi, K. Kano, S. Inokuchi, Human Exp. Toxicol. 33, 360–368 (2014).

48. M. Chaves-Ferreira, I. S. Albuquerque, D. Matak-Vinkovic, A. C. Coelho, S. M. Carvalho, L. M. Saraiva, C. C. Romão, G. J. L. Bernardes, Spontaneous CO Release from Ru^II^(CO)_2_–Protein Complexes in Aqueous Solution, Cells, and Mice. Angew. Chem. Int. Ed. 54, 1172–1175 (2015).

49. A. T. Kawaguchi, A. A. Salybekov, M. Yamano, H. Kitagishi, K. Sekine, T. Tamaki, PEGylated carboxyhemoglobin bovine (SANGUINATE) ameliorates myocardial infarction in a rat model. Artif. Organs 42, 1174–1184 (2018).

50. B. A. Springer, S. G. Sligar, J. S. Olson, G. N. Phillips, Jr., Mechanisms of ligand recognition in myoglobin. Chem. Rev. 94, 699–714 (1994).

51. J. P. Collman, J. I. Brauman, B. L. Iverson, J. L. Sessler, R. M. Morris, Q. H. Gibson, Dioxygen and carbonyl binding to iron(II) porphyrins: a comparison of the “picket fence” and “pocket” porphyrins. J. Am. Chem. Soc. 105, 3052–2064 (1983).

52. J. J. Rose, K. A. Bocian, Q. Xu, L. Wang, A. W. DeMartino, X. Chen, C. G. Corey, D. A. Guimarães, I. Azarov, X. N. Huang, Q. Tong, L. Guo, M. Nouraie, C. F. McTiernan, C. P. O’Donnell, J. Tejero, S. Shiva, M. Gladwin. A neuroglobin-based high-affinity ligand trap reverses carbon monoxide-induced mitochondrial poisoning. J. Biol. Chem. 295, 6357–6371 (2020).

53. R. C. Tuckey, H. Kamin, Kinetics of O_2_ and CO binding to adrenal cytochrome P-450scc. Effect of cholesterol, intermediates, and phosphatidylcholine vesicles. J. Biol. Chem. 258, 4232–4237 (1983).

54. E. M. Dioum, J. Rutter, J. R. Tuckerman, G. Gonzalez, M.-A. Gilles-Gonzalez, S. L. McKnight, NPAS2: A gas-responsive transcription factor. Science 298, 2385–2387 (2003).

55. I. Azarov, L. Wang, J. J. Rose, Q. Xu, X. N. Huang, A. Belanger, Y. Wang, L. Guo, C. Liu, K. B. Ucer, C. F. McTiernan, C. P. O’Donnell, S. Shiva, J. Tejero, D. B. Kim-Shapiro, M. T. Gladwin, Five-coordinate H64Q neuroglobin as a ligand-trap antidote for carbon monoxide poisoning. Sci. Transl. Med. 8, 368ra173 (2016).

56. M. Puranika, S. B. Nielsena, H. Yound, A. N. Hvitvede, J. L. Bourassaa, M. A. Casea, C. Tengrotha, G. Balakrishnana, M. V. Thorsteinssond, J. T. Groves, G. L. McLendona, G. P. Robertsd, J. S. Olson, T. G. Spiro, Dynamics of carbon monoxide binding to CooA, J. Biol. Chem. 279, 21096–21108 (2004).

57. L. Bouzhir-Sima, R. Motterlini, J. Gross, M. H. Vos, U. Liebl, Unusual dynamics of ligand binding to the heme domain of the bacterial CO sensor protein RcoM-2. J. Phys. Chem. B. 120, 10686–10694 (2016).

58. B. W. Michel, A. R. Lippert, C. J. Chang, A reaction-based fluorescent probe for selective imaging of carbon monoxide in living cells using a palladium-mediated carbonylation. J. Am. Chem. Soc. 134, 15668–15671 (2012).

59. J. Wang, J. Karpus, B. S. Zhao, Z. Luo, P. R. Chen, C. He, A selective fluorescent probe for carbon monoxide imaging in living cells. Angew. Chem. Int. Ed. 51, 9652–9656 (2012).

60. D. Madea, M. Martínek, L. Muchová, J. Váňa, L. Vítek, P. Klán, Structural modifications of nile red carbon monoxide fluorescent probe: sensing mechanism and applications. J. Org. Chem. 85, 3473–3489 (2020).

61. S. Gong, J. Hong, E. Zhou, G. Feng, A near-infrared fluorescent probe for imaging endogenous carbon monoxide in living systems with a large Stokes shift. Talanta 201, 40–45 (2019).

62. K. Liu, X. Kong, Y. Ma, W. Lin, Rational design of a robust fluorescent probe for the detection of endogenous carbon monoxide in living zebrafish embryos and mouse tissue. Angew. Chem. Int. Ed. 56, 13489–13492 (2017).

63. Y. Li, X. Wang, J. Yang, X. Xie, M. Li, J. Niu, L. Tong, B. Tang, Fluorescent probe based on azobenzene-cyclopalladium for the selective imaging of endogenous carbon monoxide under hypoxia conditions. Anal. Chem. 88, 11154–11159 (2016).

64. Y. Morimoto, W. Durante, D. G. Lancaster, J. Klattenhoff, F. K. Tittel, Real-time measurements of endogenous CO production from vascular cells using an ultrasensitive laser sensor. Am. J. Physiol. Heart Circ. Physiol. 280, H483–H488 (2001).

65. G. S. Marks, H. J. Vreman, B. E. McLaughlin, J. F. Brien, K. Nakatsu, Measurement of endogenous carbon monoxide formation in biological systems. Antioxid. Redox Signal. 4, 271–277 (2002).

66. H. M. Deloar, H. Watabe, T. Nakamura, Y. Narita, A. Yamadera, T. Fujiwara, M. Itoh, Internal dose estimation including the nasal cavity and major airway for continuous inhalation of C^15^O_2_, ^15^O_2_ and C^15^O using the thermoluminescent dosimeter method. J. Nucl. Med. 38, 1603–1613 (1997).

67. S. Oh, S.-C. Choi, Acute carbon monoxide poisoning and delayed neurological sequelae: a potential neuroprotection bundle therapy. Neural. Regen. Res. 10, 36–38 (2015).

68. S. D. Brown, C. A. Piantadosi, In vivo binding of carbon monoxide to cytochrome c oxidase in rat brain. J. Appl. Physiol. 68, 604–610 (1985).

69. J. Zhang, C. A. Piantadosi, Mitochondrial oxidative stress after carbon monoxide hypoxia in the rat brain. J. Clin. Invest. 90, 1193–1199 (1992).

70. S. D. Brown, C. A. Piantadosi, Recovery of energy metabolism in rat brain after carbon monoxide hypoxia. J. Clin. Invest. 89, 666–672 (1992).

