## Supplementary Information for "Highly sensitive quantification of carbon monoxide (CO) *in vivo* reveals a protective role of circulating hemoglobin in CO intoxication"

**This PDF file includes:**

Materials and Methods
References in Supplementary Information
Table S1. Molar absorption coefficients of hemoCD1 used in this study.
Fig. S1. Stabilities of CO-hemoCD1 and CO-Hb upon $O_2$ bubbling.
Fig. S2. Stabilities of CO-hemoCD1 and CO-Hb upon $N_2$ bubbling.
Fig. S3. Stabilities of CO-hemoCD1 and CO-Hb upon vacuuming.
Fig. S4. Stabilities of CO-hemoCD1 and CO-Hb against oxidation.
Fig. S5. Stabilities of CO-, deoxy-, and oxy-hemoCD1 in the solution containing biocomponents from cell lysates.
Fig. S6. Ligand exchange of CO-Hb and oxy-hemoCD1.
Fig. S7. Stabilities of CO-hemoCD1 and CO-Hb against NO.
Fig. S8. Effect of Py3CD and hemoCD1 on cell viability.
Fig. S9. Quantification of endogenous CO in tissues.
Fig. S10. Plots of the wet weight of tissues versus the amount of CO detected by the hemoCD1 assay.
Fig. S11. Effects of external reactive species, pH, and light on quantification of endogenous CO in the liver and brain tissues by hemoCD1.
Fig. S12. Comparison between the hemoCD1 assay and the gas chromatography method for CO quantification in tissues.
Fig. S13. Schematic representation of the experimental setup for CO gas inhalation in rats.
Fig. S14. *In vitro* experiments to demonstrate CO transfer from tissues to Hb.
Fig. S15. UV-vis spectra of rat urine collected after i.v. infusion of oxy-hemoCD1 to CO-treated rats.
Fig. S16. Effect of air/$O_2$ ventilation in combination with hemoCD1 on CO distribution in tissues after CO inhalation *in vivo*.
Fig. S17. Effect of $O_2$ ventilation in combination with hemoCD1 on CO distribution in tissues after CO inhalation *in vivo*.

### Materials and Methods

#### Preparation of ferrous hemoCD1

5,10,15,20-Tetrakis(4-sulfonatopheyl)porphinatoiron(III) ($Fe^{III}$TPPS) and Py3CD were synthesized in our laboratory (*S1,S2*). Stock solutions of hemoCD1 used for CO quantification in tissues were prepared as follows. $Fe^{III}$TPPS (1.10 mg, 1.0 μmol) and Py3CD (3.53 mg, 1.2 μmol) were dissolved in PBS (1 mL) to yield a solution of met-hemoCD1 (1 mM). The stock solution of met-hemoCD1 (1–5 μL) was appropriately diluted with PBS to prepare the solution of met-hemoCD1 (2–10 μM, 0.5 mL) for CO quantification in tissues. Ferrous deoxy-hemoCD1 was obtained by adding an excess of $Na_2S_2O_4$ (*ca.* 1–2 mg) to the met-hemoCD1 solution (2–10 μM, 0.5 mL).

The solution of oxy-hemoCD1 used as a CO removal agent to be injected in rats was prepared as follows. $Fe^{III}$TPPS (6.60 mg, 6.0 μmol) and Py3CD (21.15 mg, 7.2 μmol) were dissolved in PBS (2 mL) to yield a solution of met-hemoCD1 (3 mM). $Na_2S_2O_4$ (*ca.* 10–20 mg) was then added to reduce met-hemoCD1 to ferrous deoxy-hemoCD1. Excess $Na_2S_2O_4$ was removed by passing the deoxy-hemoCD1 solution through a HiTrap desalting column (Sephadex G25, GE Healthcare Life Sciences). During the filtration process, deoxy-hemoCD1 was converted to oxy-hemoCD1 by capturing atmospheric $O_2$. The concentration of oxy-hemoCD1 was determined from its absorption coefficient $\varepsilon_{422}{}^{oxy} = 1.64 \times 10^5\ M^{-1}\ cm^{-1}$ (*S3*).

#### Animals

Five weeks old Lewis and Sprague-Dawley rats were purchased from Charles River Co. Ltd., (Yokohama, Japan). Before undergoing any experiment, animals were acclimatized for a week under air conditioning at 26 ± 0.5°C with access to water and food ad libitum.

#### Animal preparation

All experiments were approved by the Institutional Review Board of Tokai University and the Guidelines for Animal Experiments of Doshisha University. Animals received humane care as required by the institutional guidelines for animal care and treatment in experimental investigations according to the Guide for the Care and Use of Laboratory Animals (Institute of Laboratory Animal Resources, 1996). Rats were anesthetized with 3% sevoflurane, orally intubated and ventilated at 15 mL/kg of tidal volume with ambient air at a ventilation rate of 55 per minute with no end-expiratory pressure by a ventilator (Rodent Ventilator, Ugo Basile). An indwelling catheter was placed in the tail vein and saline was administered at 3 mL/h. The body temperature was monitored with a rectal probe and maintained at 36.5 ± 0.5°C with a water blanket (MEDI-Therm II, Gaymer Industries Inc.). The chest was opened at the $4^{th}$ intercostal space, the dose of sevoflurane was reduced to 2%, and rats were anesthetized with an intraperitoneal injection of pentobarbital (PB). After 5 min, sevoflurane anesthesia was removed and rats were ready for the experiments with CO gas inhalation (see below).

**Blood CO-Hb measurements**

Rat arterial and venous blood samples (0.1 mL) were regularly collected from left and right ventricles (LV and RV). Blood samples were immediately analyzed using a blood gas analyzer, ABL825 (Radiometer Co. Ltd.), which measures CO-Hb based on a 128 wavelengths spectrometer with a measuring range from 478 to 672 nm. As shown in Fig. 4–6, CO-Hb in arterial blood was always 1-2% higher than that in the venous sample, the difference being consistent with a previous report (*S4*).

**Preparation of tissue samples**

Tissue specimens from liver, lung, cerebrum, cerebellum, heart (myocardium), and skeletal muscle were collected immediately after the chest was opened. To remove blood from the organs, saline was flushed before collecting the tissues as follows: the blood circulation was perfused with saline (50 mL) through the pulmonary artery and then puncturing the left ventricle with a 16G needle (TERUMO). Additional saline (150 mL) was then injected at 0.25 mL/sec using a peristaltic pump. The flushed tissues were collected and immediately frozen in liquid nitrogen and stored at –80°C prior to CO measurements.

**Quantification of CO in tissues by the hemoCD1 assay**

Tissue samples (5-20 mg) harvested from rat organs were weighed and homogenized by Power MasherII (nippi) in PBS (0.5 mL). After homogenization, deoxy-hemoCD1 (2–10 μM) with $Na_2S_2O_4$ (*ca*. 1–2 mg) in PBS (0.5 mL) was added to the tissue homogenates, which were then disrupted by sonication (time: 10 s × 2, on ice, amplitude: 15; QSONICA). Samples were centrifuged (14000 G, 15 min) and the supernatants filtered (DISMIC 13CP, 0.45 μm pore; ADVANTEC). The filtrates were treated with $Na_2S_2O_4$ (*ca*. 1–2 mg) before measurements by UV-vis absorption spectroscopy (NanoPhotometer® C40, Implen). The concentration of total hemoCD1 ($C_{total}$) was determined from the absorbance at 427 nm by eq 1 as follows:

$$A_{427\ nm} = \varepsilon_{427} \cdot C_{total} \cdot l \qquad (1)$$

where $\varepsilon_{427}$ is the molar extinction coefficient at 427 nm ($1.95 \times 10^5$ $M^{-1}cm^{-1}$), at which the wavelength of the isosbestic point of deoxy- and CO-hemoCD1. The $l$ is the optical path length (1.0 cm). The resulting hemoCD1 solution contains CO-hemoCD1 ($C_{co}$) and deoxy-hemoCD1 ($C_{deoxy}$) (eq2):

$$C_{total} = C_{co} + C_{deoxy} \tag{2}$$

The ratio of CO-hemoCD1 in total hemoCD1 ($R_{CO}$) is represented as eq 3:

$$R_{CO} = C_{co} / C_{total} \tag{3}$$

The absorbances at 422 nm and 434 nm are defined as eqs 4 and 5:

$$A_{422} = \varepsilon_{deoxy}{}^{422} \cdot C_{deoxy} + \varepsilon_{co}{}^{422} \cdot C_{co} \tag{4}$$

$$A_{434} = \varepsilon_{deoxy}{}^{434} \cdot C_{deoxy} + \varepsilon_{co}{}^{434} \cdot C_{co} \tag{5}$$

Therefore, the ratio of $A_{422}/A_{434}$ is represented as eq 6:

$$A_{422}/A_{434} = \frac{\varepsilon_{deoxy}{}^{422} \cdot C_{deoxy} + \varepsilon_{co}{}^{422} \cdot C_{co}}{\varepsilon_{deoxy}{}^{434} \cdot C_{deoxy} + \varepsilon_{co}{}^{434} \cdot C_{co}} \tag{6}$$

From eqs 2 to 6, $R_{CO}$ is represented as eq 7:

$$R_{CO} = \frac{\varepsilon_{deoxy}{}^{422} - A_{422}/A_{434} \cdot \varepsilon_{deoxy}{}^{434}}{A_{422}/A_{434}\,(\varepsilon_{co}{}^{434} - \varepsilon_{deoxy}{}^{434}) - \varepsilon_{co}{}^{422} + \varepsilon_{deoxy}{}^{422}} \tag{7}$$

where $\varepsilon_{co}{}^{422} = 3.71 \times 10^5$ $M^{-1}cm^{-1}$, $\varepsilon_{co}{}^{434} = 6.75 \times 10^4$ $M^{-1}cm^{-1}$, $\varepsilon_{deoxy}{}^{422} = 1.52 \times 10^5$ $M^{-1}cm^{-1}$, and $\varepsilon_{deoxy}{}^{434} = 2.13 \times 10^5$ $M^{-1}cm^{-1}$ are used for calculation. The moles of CO contained as CO-hemoCD1 in solution ($M_{CO}$) are determined by eq 8:

$$M_{CO}\ (\text{mol}) = R_{CO} \cdot C_{total} \cdot V \tag{8}$$

where $V$ is the volume of the solution ($1 \times 10^{-3}$ L). The relation between the $R_{CO}$ and $A_{422}/A_{434}$ values in eq 7 is nonlinear, and thus reproducibility of $R_{CO}$ would become low when $A_{422}/A_{434}$ below 0.8 ($R_{CO}$ = 0.055) or over 3.0 ($R_{CO}$ = 0.743). For accurate quantification, the $A_{422}/A_{434}$ values were thus adjusted between 0.8 to 3.0 by slightly varying the initial amount of tissues or hemoCD1 added to the samples.

**Protocol for exposure of tissues to CO *ex vivo***

Tissue samples were placed in a 5 mL 25G syringe (TERUMO) sealed with a septum cap (natural rubber, for a 7 mm tube). The atmosphere in the syringe was replaced with pure CO gas (4 mL) (Sumitomo Seika) through inserting a 25G needle (TERUMO) into the septum cap. Tissue samples were then incubated at 4°C under a CO atmosphere. After 1 h, the syringe was opened under air atmosphere, samples were taken out from the syringe, and the amount of CO was measured by the hemoCD1 assay.

**Quantification of CO in tissues by gas chromatography**

The amount of CO in the tissue samples was measured by gas chromatography according to the reported method (*S5*). Tissues (100 mg) were homogenized (Power MasherII, nippi) in PBS (0.3 mL) and then disrupted by sonication (time: 10 s × 2, on ice, amplitude: 15; QSONICA). The solution and a 5 mmφ glass ball were placed in a 5 mL 25G syringe (TERUMO) sealed with a septum cap (natural rubber, for a 7 mm tube). The atmosphere in the syringe (1 mL) was carefully replaced with He gas through inserting a 25G needle (TERUMO) into the septum cap. Three drops of 30% sulfosalicylic acid (SSA) (Wako) were injected through the septum into the syringe. The mixture was then mixed well using a 5 mmφ glass ball. Methane gas (50 μL) was added to the head-space atmosphere through the septum in the syringe as an internal standard. CO liberated from the tissues into the gas phase (0.5 mL) was analyzed by gas chromatography (GC, Shimadzu GC-2014, Shimadzu). The GC conditions were as follows: detector = TCD; column = SHINCARBON: carrier gas = He; column temperature = 40°C; temperature at vaporizing chamber and detector = 120 °C; flow rate = 50 mL/min.

**Exposure to CO gas inhalation *in vivo***

The experimental setup for CO gas inhalation *in vivo* is shown in Figure S11. After anesthesia by an intraperitoneal (i.p.) injection of pentobarbital (PB), rats were exposed to 400 ppm CO gas equilibrated in air (flow: 1 L/min, TOMOE SHOKAI Co., LTD.) through oral intubation and ventilation (Rodent Ventilator, Ugo Basile) for 5, 10 or 20 min. The concentration of the inhaled CO gas was monitored by a CO detector (COSMOS XC2200). The exhaled gas was diluted with air before exiting and the atmosphere was monitored during the entire experiment. During CO exposure, arterial and venous blood from right and left ventricles (RV and LV, 0.1 mL each) were collected and analyzed using the blood gas analyzer as described above. At each time point, organs were flushed as described above and tissue samples collected and frozen were finally stored at –80°C for further CO measurements.

**Protocols of air/$O_2$ ventilations after exposure to CO *in vivo***

The experimental setup was the same as shown in Figure S11. Anesthetized rats were exposed to CO gas (400 ppm) for 5 min as described above, and then breathing gas was changed to either air or pure $O_2$ (TOMOE SHOKAI Co., LTD.). Arterial and venous blood samples (0.1 mL) were collected for blood gas analyses and tissues were harvested at different time points. Tissues were frozen in liquid nitrogen and stored at –80°C for further CO measurements.

**Administration of oxy-hemoCD1 to rats exposed to CO**

The solution of oxy-hemoCD1 (1.4–3.0 mM, 2.5 mL in PBS) was freshly prepared before each experiment to avoid autoxidation. Before the experiments, we confirmed that in the blood gas analysis (ABL825, Radiometer Co. Ltd.) at 128 different wavelengths the Hb spectrum was unaffected by the oxy-hemoCD1 administration because the injected amount of oxy-hemoCD1 was much less (*ca.* 1/100) than Hb in blood.

In the case of the experiments conducted according to protocol ***I***, the solution of oxy-hemoCD1 (1.4–3.0 mM, 2.5 mL in PBS) was first prepared in a syringe and placed in a syringe pump (TERUMO) ready to be injected. Five minutes after exposure of rats to CO gas (400 ppm), the inhaled gas was switched to air and 1 mL of oxy-hemoCD1 was quickly infused from the tail vein. The rest of the solution of oxy-hemoCD1 (1.5 mL) was then infused through the same vein at a rate of 4.5 mL/h (the total infusion time: 30 min). At the end of infusion, rats were kept on air ventilation for an additional 30 min. Arterial and venous blood samples were collected at 0, 5, 15, 35, and 65 min time points ($t_0$, $t_5$, $t_{15}$, $t_{35}$, $t_{65}$). At the same time points, the urine was collected and analyzed by UV-vis absorption spectroscopy (NanoPhotometer® C40, Implen). At each time point, the chest was opened, tissues were flushed using saline (200 mL), collected and frozen in liquid nitrogen. Samples were finally stored at –80°C for further CO measurements.

In the case of the experiments carried out according to protocol ***II***, the procedure was the same as in protocol ***I*** except that after exposure of rats to CO, the inhaled gas was switched to pure $O_2$ and 1 mL of oxy-hemoCD1 was quickly infused intravenously. The rest of the solution of oxy-hemoCD1 (1.5 mL) was then infused in the same vein at a rate of 9.0 mL/h (the total infusion time: 15 min). After the end of infusion, the rats were kept on pure $O_2$ ventilation for an additional 45 min. Arterial and venous blood samples and tissues were collected using the same procedures as in protocol ***I***.

In the case of the experiments conducted according to protocol ***III***, rats were treated with CO gas as in the other protocols, then the gas was switched to pure $O_2$. After 30 min, a solution of oxy-hemoCD1 (1 mL) was quickly infused from the tail vein and the rest (1.5 mL) was infused to the same vein at a rate of 9.0 mL/h (the total infusion time: 15 min). At the end of infusion, rats were kept on $O_2$ ventilation for an additional 15 min. Arterial and venous blood samples and tissues were collected using the same procedures as in protocol ***I***.

In the case of the experiments conducted according to protocol ***IV*** and ***V*** (Figure 6B), rats were treated with 400 ppm CO gas for 80 min, then the gas was switched to pure $O_2$. In case of 80 min inhalation, 3.0 mM oxy-hemoCD solution (2.5 mL) was used for intravenous injection. Subsequent procedures were the same as protocols ***II*** and ***III***, respectively.

**Cell cultures**

The human hepatoma cell line (HepG2) was obtained from RIKEN Cell Bank and cultured in Dulbecco's modified Eagle's medium (DMEM, GIBCO) supplemented with 10% fetal bovine serum (FBS, Invitrogen GIBCO, heat inactivated at 56 °C before use) and 1% penicillin/streptomycin/amphotericin B (Wako) at 37°C in a humidified atmosphere in the presence of 5% $CO_2$. Human met-Hb was purchased from Sigma-Aldrich. Ferrous Hb was prepared using $Na_2S_2O_4$ as a reducing agent, followed by purification using a HiTrap desalting column, Sephadex G25, GE Healthcare Life Sciences. The concentration of Hb was determined using reported absorption coefficients (*S6*).

***In vitro* experiments to demonstrate CO transfer from cells to Hb**

HepG2 ($1 \times 10^6$) cells were treated with CORM401-E (25 μM in D-10) for 2 h at 37°C. The medium was then removed, cells were washed with PBS, and then incubated with oxy-Hb (15 μM in D-10) for 1 h. Cells were washed with PBS, collected by scraping and the suspension was sonicated. The amounts of CO contained in the cells were then quantified by the hemoCD1 assay.

**Effect of hemoCD1 and Py3CD on cell viability**

Cell viability was assessed by the MTT assay. Briefly, HepG2 cells (*ca*. $10^5$ cells) in D-10 (100 μL) were seeded into each well of 96-well plates and incubated at 37 °C in a humidified $CO_2$ atmosphere (5%) for 24 h. The medium was replaced with fresh medium

(D-10) containing Py3CD (20, 50, 100, 150, and 250 μM ) or met-hemoCD1 (20, 50, 100, 150, and 250 μM) and treated for 3 h. MTT (Sigma) was dissolved in PBS at a concentration of 5 mg/mL. After 3 h, 100 μL of medium was removed and 10 μL of the MTT solution with 90 μL D-10 was added to each well. After 3 h incubation at 37°C, 100 μL of medium was removed and mixed with 100 μL DMSO (Wako) before absorbance at 570 nm was determined on a Multiskan™ FC Microplate Photometer (Thermo Fisher Scientific).

**Statistical analysis.**

All data represent the means ± standard error from at least three different experiments and were analyzed by unpaired Student's *t* tests. Differences with *P* values of less than 0.05 were considered significant.

**References in Experimental Section**

S1. E. B. Fleischer, J. M. Palmer, T. S. Srivastava, A. Chatterjee, Thermodynamic and kinetic properties of an iron-porphyrin system. *J. Am. Chem. Soc.* **93**, 3162–3167 (1971).

S2. K. Kano, H. Kitagishi, M. Kodera, S. Hirota, Dioxygen binding to a simple myoglobin model in aqueous solution. *Angew. Chem. Int. Ed.* **44**, 435–438 (2005).

S3. S. Minegishi, A. Yumura, H. Miyoshi, S. Negi, S. Taketani, R. Motterlini, R. Foresti, K. Kano, H. Kitagishi, Detection and removal of endogenous carbon monoxide by selective and cell-permeable hemoprotein-model complexes. *J. Am. Chem. Soc.*, **139**, 5984–5991 (2017).

S4. M. Touger, E. J. Gallagher, J. Tyrell, Relationship between venous and arterial carboxyhemoglobin levels in patients with suspected carbon monoxide poisoning. *Ann. Emerg. Med.* **25**, 481–483 (1995).

S5. H. J. Vreman, R. J. Wong, T. Kadotani, T. K. Stevenson, Determination of carbon monoxide (CO) in rodent tissue: Effect of heme administration and environmental CO exposure. *Anal. Biochem.* **341**, 280–289 (2005).

S6. E. Antonini, M. Brunori, "Hemoglobin and myoglobin in their reactions with ligands" in Frontiers in Biology, A. Neuberger, E. L. Tatum, Eds. (Elsevier: Amsterdam, 1971), Chapter 13.

**Table S1.** Molar absorption coefficients ($\varepsilon$ / $M^{-1}$ $cm^{-1}$) of hemoCD1 used in this study.

| | 422 nm | 427 nm | 434 nm |
|---|---|---|---|
| Deoxy-hemoCD1 | 152 000[b] | 195 000[b] | 213 000[a] |
| CO-hemoCD1 | 371 000[a] | 195 000[b] | 67 500[b] |

[a]Ref (*31,39*). [b]This work.

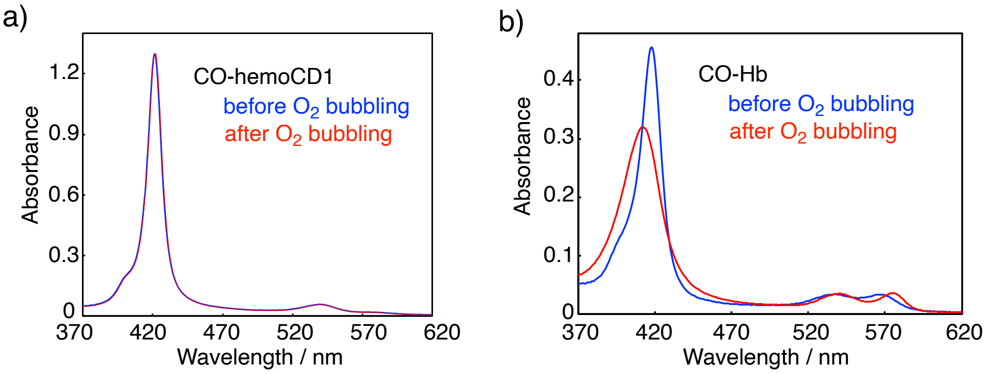


**Fig. S1. Stabilities of CO-hemoCD1 and CO-Hb upon $O_2$ bubbling.** (a,b) UV-vis spectra of CO-hemoCD1 (a) and CO-Hb (b) in PBS at pH 7.4 and 25°C before (blue) and after (red) bubbling $O_2$ into the solutions for 5 min. CO-Hb was converted to its $O_2$-bound form (oxy-Hb, red line) whereas CO-hemoCD1 was not.

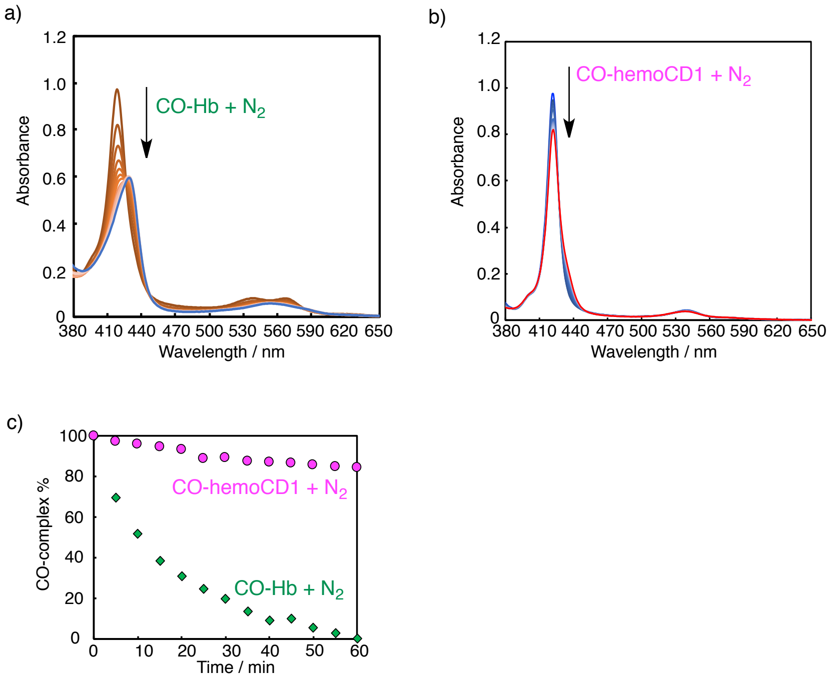


**Fig. S2. Stabilities of CO-hemoCD1 and CO-Hb upon $N_2$ bubbling.** (a,b) UV-vis spectral changes of CO-Hb (a) and CO-hemoCD1 (b) in phosphate buffer saline (PBS) at pH 7.4 and 25°C after $N_2$ bubbling. (c) The plot of the residual CO-complexes (%) of Hb and hemoCD1 during $N_2$ bubbling. The data indicate that CO-Hb was converted to its deoxy-form by reducing CO partial pressure, while CO-hemoCD1 was hardly affected due to its high CO binding affinity and stability.

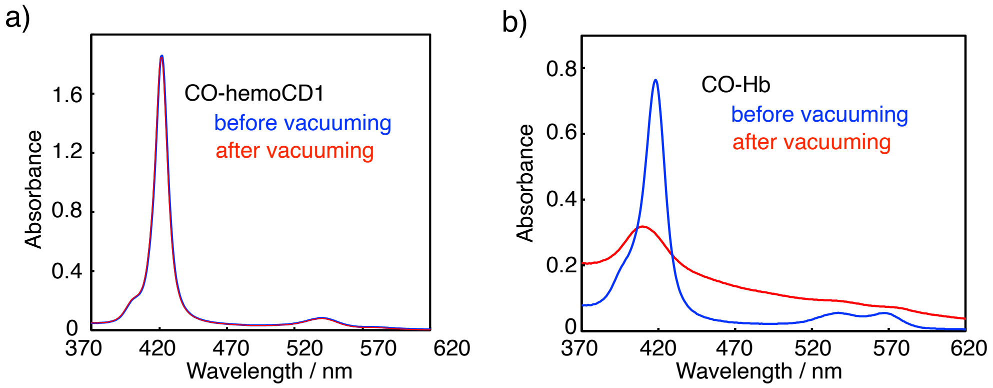


**Fig. S3. Stabilities of CO-hemoCD1 and CO-Hb upon vacuuming.** (a,b) UV-vis spectra of CO-hemoCD1 (a) and CO-Hb (b) before (blue) and after (red) vacuuming under 10 Torr for 2 h followed by re-solubilization of the residues. CO-hemoCD1 was stable while CO-Hb decomposed after this treatment.

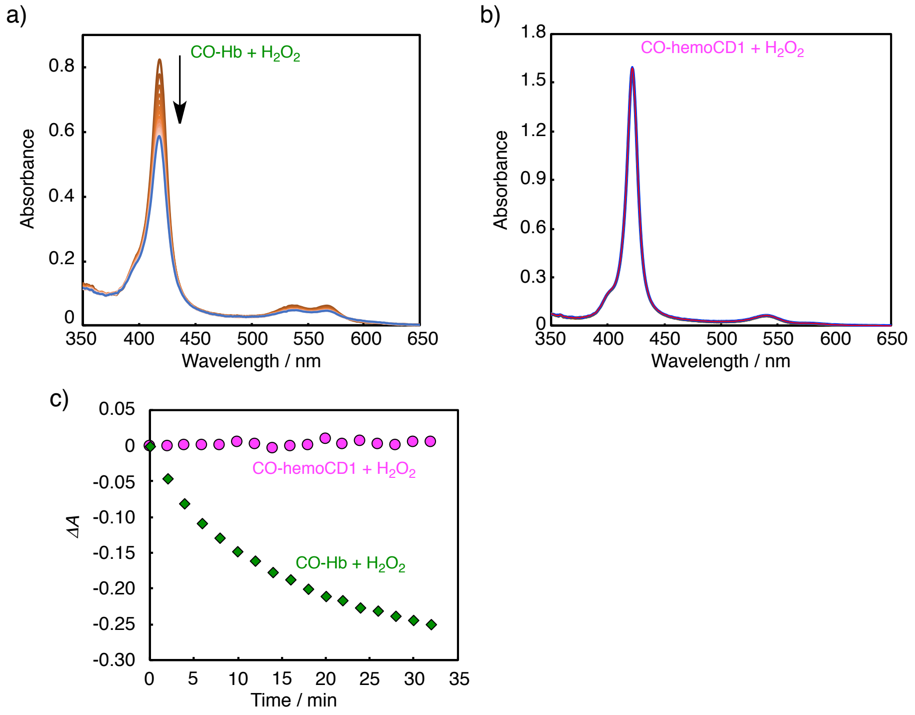


**Fig. S4. Stabilities of CO-hemoCD1 and CO-Hb against oxidation.** (a,b) UV-vis spectra of CO-Hb (a) and CO-hemoCD1 (b) in phosphate buffer saline (PBS) at pH 7.4 and 25°C after the addition of 20 eq of $H_2O_2$ at 2 min recording time intervals. (c) The plot of the changes in absorbances at $\lambda_{max}$ of CO-Hb (418 nm) and CO-hemoCD1 (422 nm). The data indicate that CO-Hb was gradually oxidized and decomposed by $H_2O_2$, while CO-hemoCD1 was resistant to oxidation by $H_2O_2$.

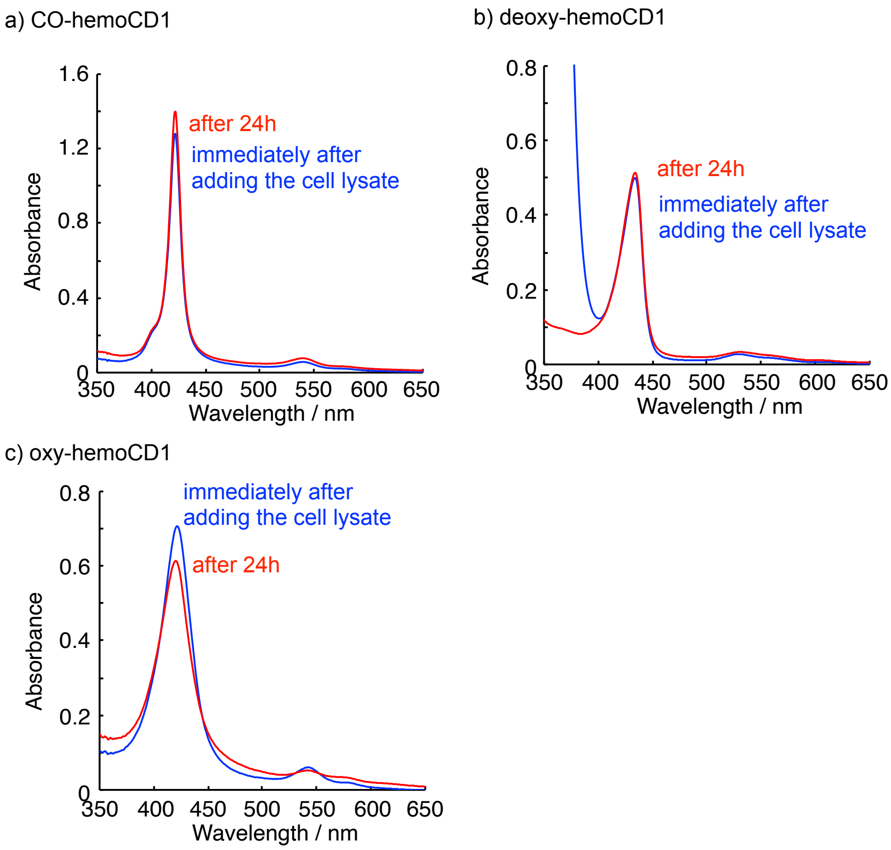


**Fig. S5. Stabilities of CO-, deoxy-hemoCD1 and oxy-hemoCD1 in the solution containing biocomponents from cell lysates ($10^6$ cells).** The spectra were measured at 25°C immediately after adding the cell lysate and after 24 h. Deoxy- and CO-hemoCD1 were quite stable over 24 h while oxy-hemoCD1 was gradually autoxidized to ferric met-form in the same manner as in the absence of cell lysates.

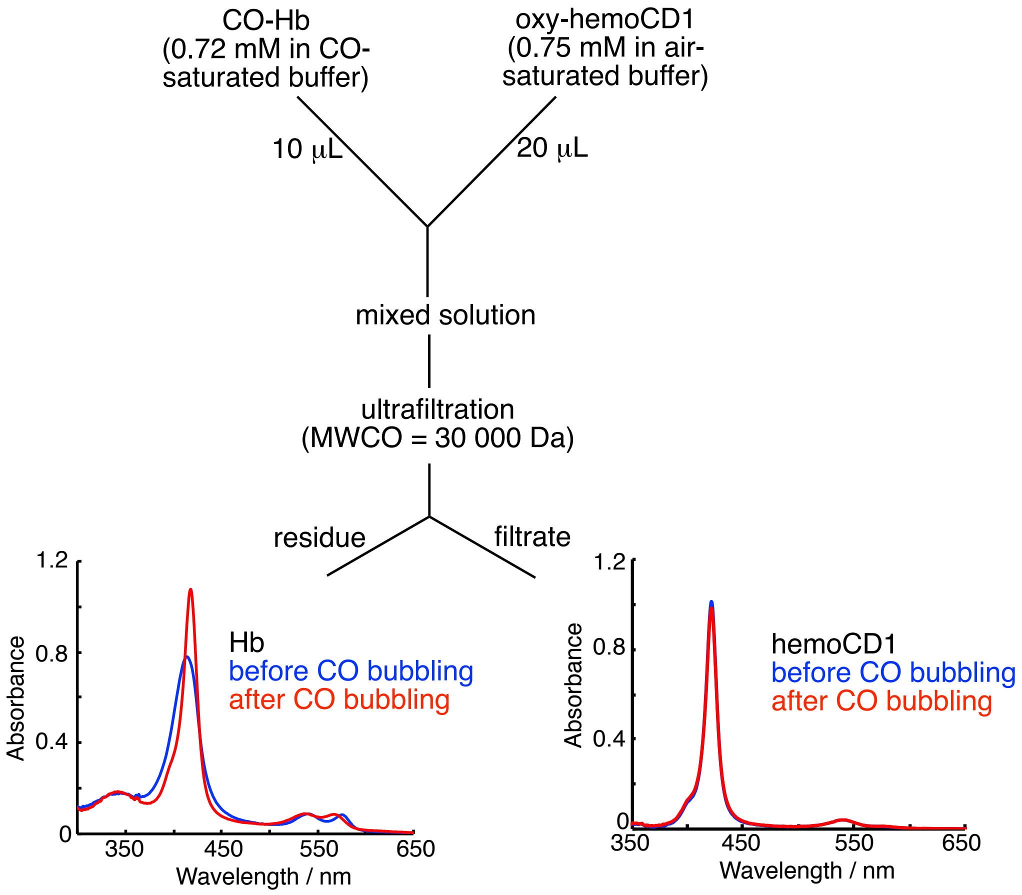


**Fig. S6**. **An experiment for ligand exchange of CO-Hb and oxy-hemoCD1 using an ultrafilter unit.** Two solutions contained CO-Hb (0.72 mM in CO-saturated buffer) and oxy-hemoCD1 (0.75 mM in air-saturated buffer) were mixed then filtered using an ultrafilter with molecular weight cut off = 30 000 Da (Amicon Ultra). The resulting residue and filtrate were appropriately diluted with PBS for UV-vis measurement. CO gas was bubbled to each solution after the measurement. The spectral characteristics indicate the residue containing significant amount of oxy-Hb and the filtrate mostly containing CO-hemoCD1, as a result of quantitative ligand exchange between each other.

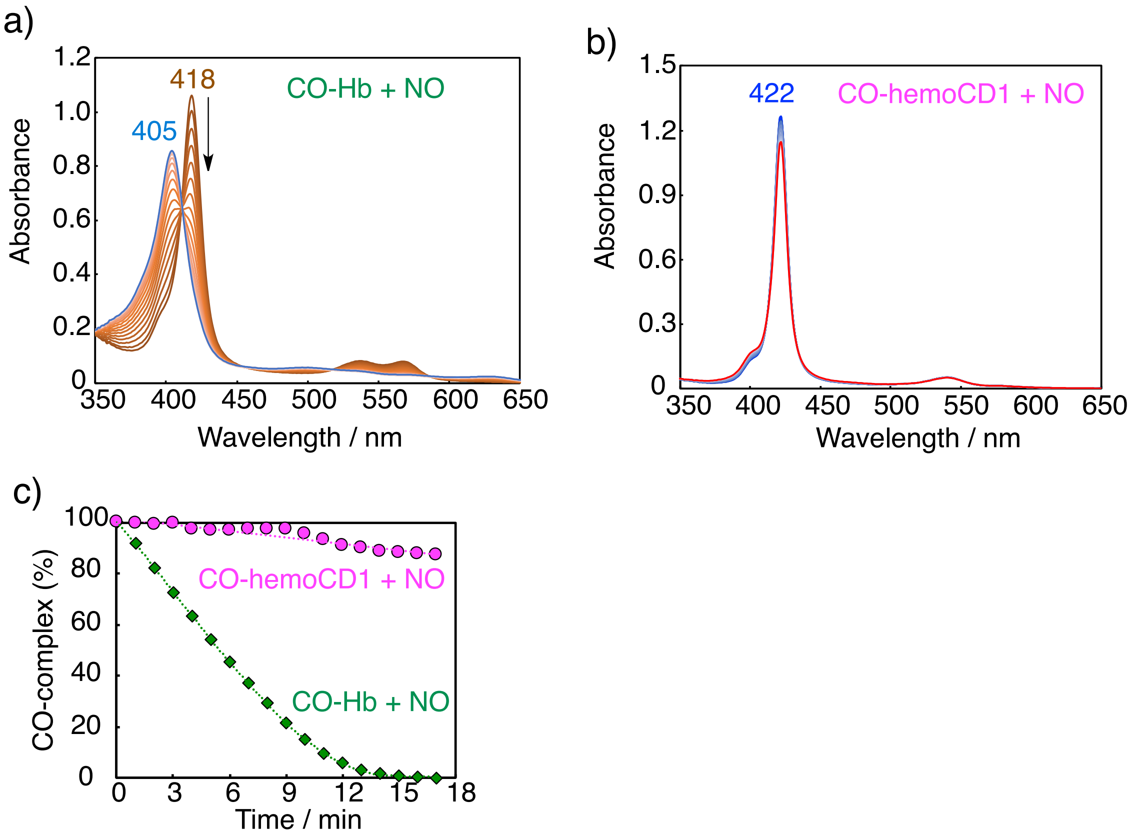


**Fig. S7**. **Stabilities of the CO-complexes of Hb and hemoCD1 against NO.** (a,b) UV-vis spectra of CO-Hb (a) and CO-hemoCD1 (b) after addition of NO (2.5 eq). NO was added as NOC (1-hydroxy-2-oxo-3-(3-aminopropyl)-3-isopropyl-1-triazene), a NO donor purchased from Dojin Chemicals. (c) The plot of the residual CO-complexes (%) of Hb and hemoCD1 during the reaction with NO. The data indicate that CO-Hb was smoothly converted to its met-form by the addition of NO, as we reported in a previous paper (H. Kitagishi et al., *J. Am. Chem. Soc.* **2016**, *138*, 5417, Supporting Information). In the case of CO-hemoCD1, its spectra were almost unchanged after reaction with NO due to the high stability of the CO-complex. This indicates that the CO binding affinity to hemoCD1 is much higher than that for NO. In addition, as we discussed in *J. Am. Chem. Soc.* **2017**, *139*, 5984 (Supporting Information), hemoCD1 cannot scavenge NO from NO-Hb due to its lower NO affinity compared to Hb. This indicates that NO does not affect the CO-scavenging ability of hemoCD1.

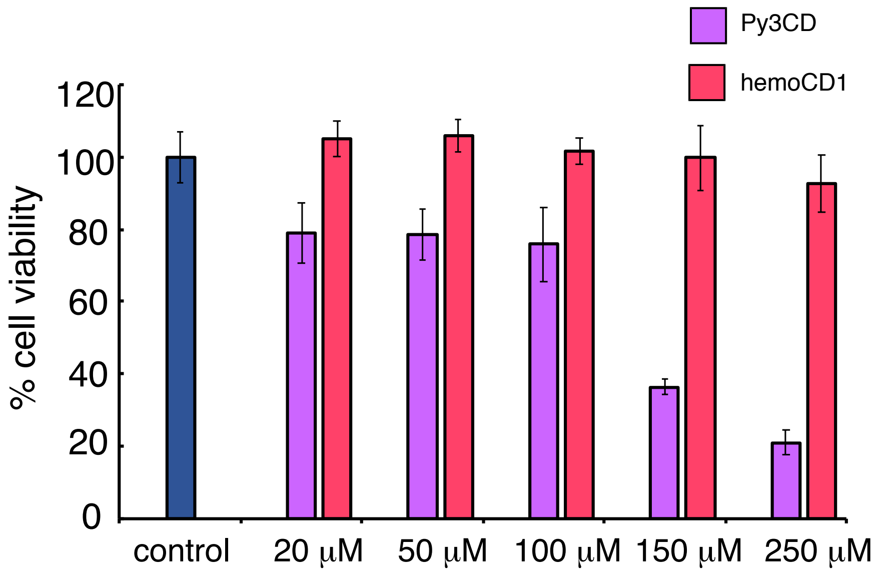


**Fig. S8**. **Effect of Py3CD and hemoCD1 on cell viability.** Hepatocytes were incubated with different concentrations of Py3CD or met-hemoCD1 for 3 h. Cell viability was measured using an MTT assay. Each bar represents the mean ± SE ($n$ = 6).

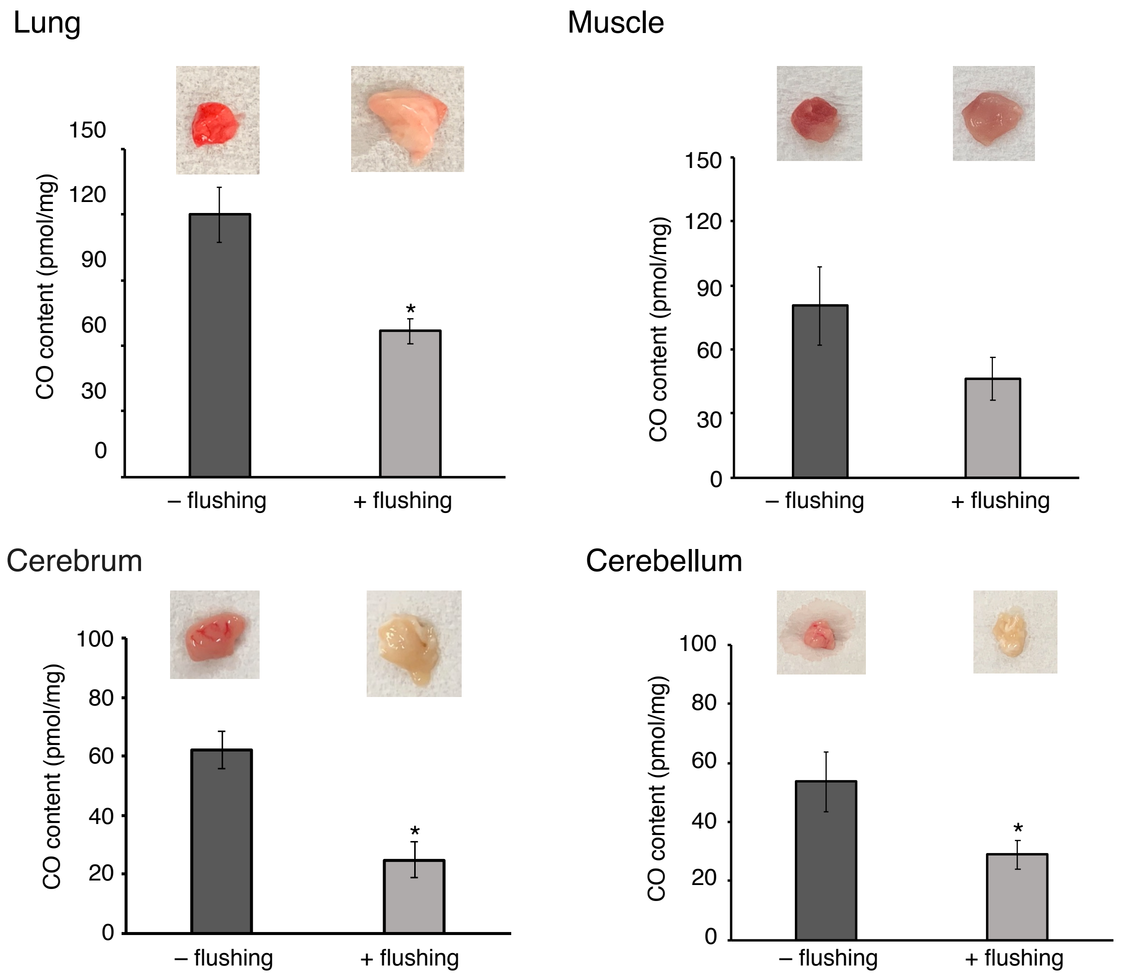


**Fig. S9**. **Quantification of endogenous CO in tissues.** The amounts of CO contained in the lung, muscle, cerebrum, and cerebellum without (–) and with flushing (+) the organs with saline (200 mL) to remove residual blood. The amounts of CO were quantified by the hemoCD1 assay. Each bar represents the means ± SE ($n$ = 3–6 rats per group). *$p < 0.05$ vs –flushing.

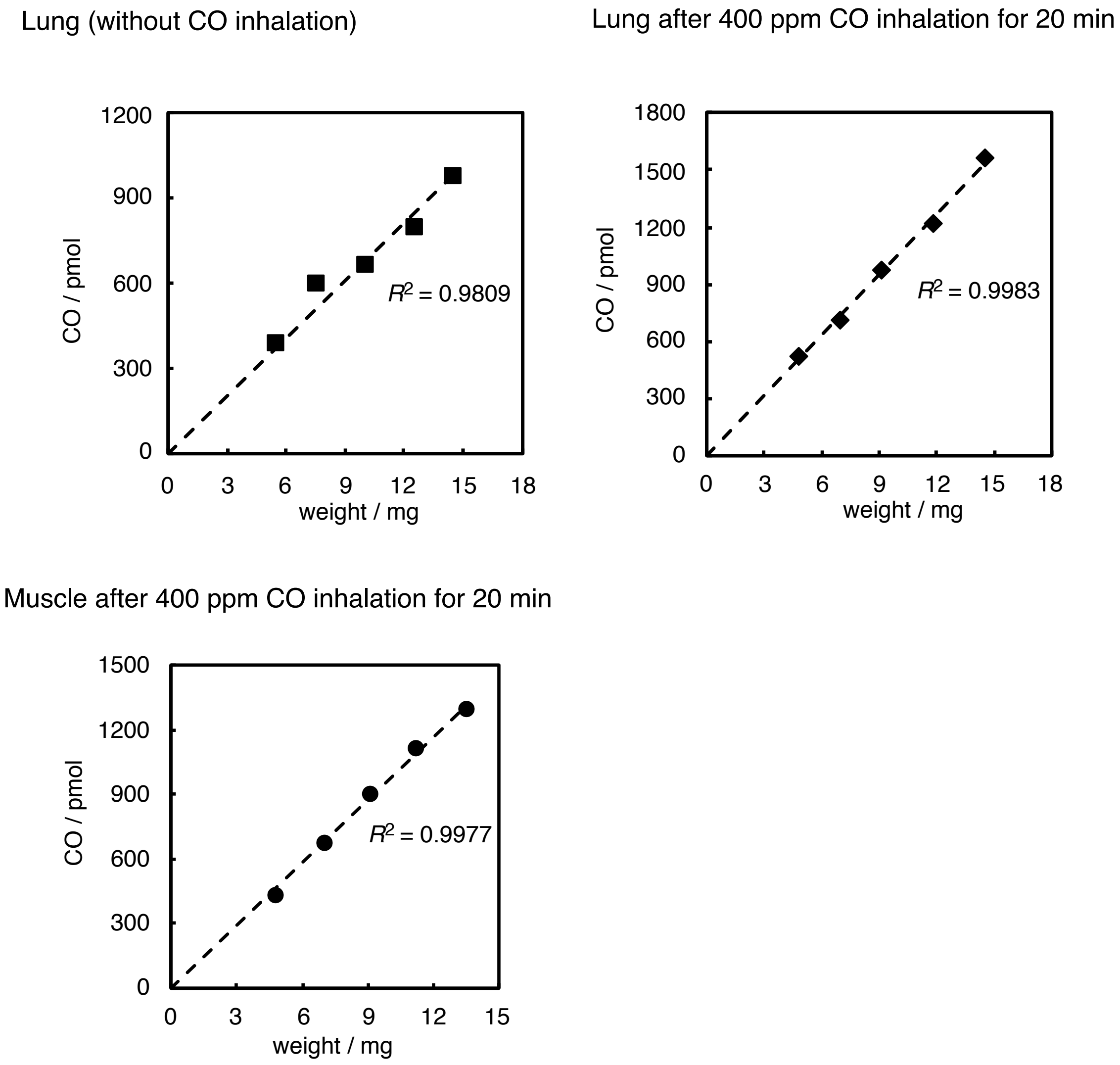


**Fig. S10**. **Plots of the wet weight of tissues (lung and muscle, with and without CO inhalation) versus the amount of CO detected by the hemoCD1 assay.** The linearity ensures accuracy of the CO quantification assay.

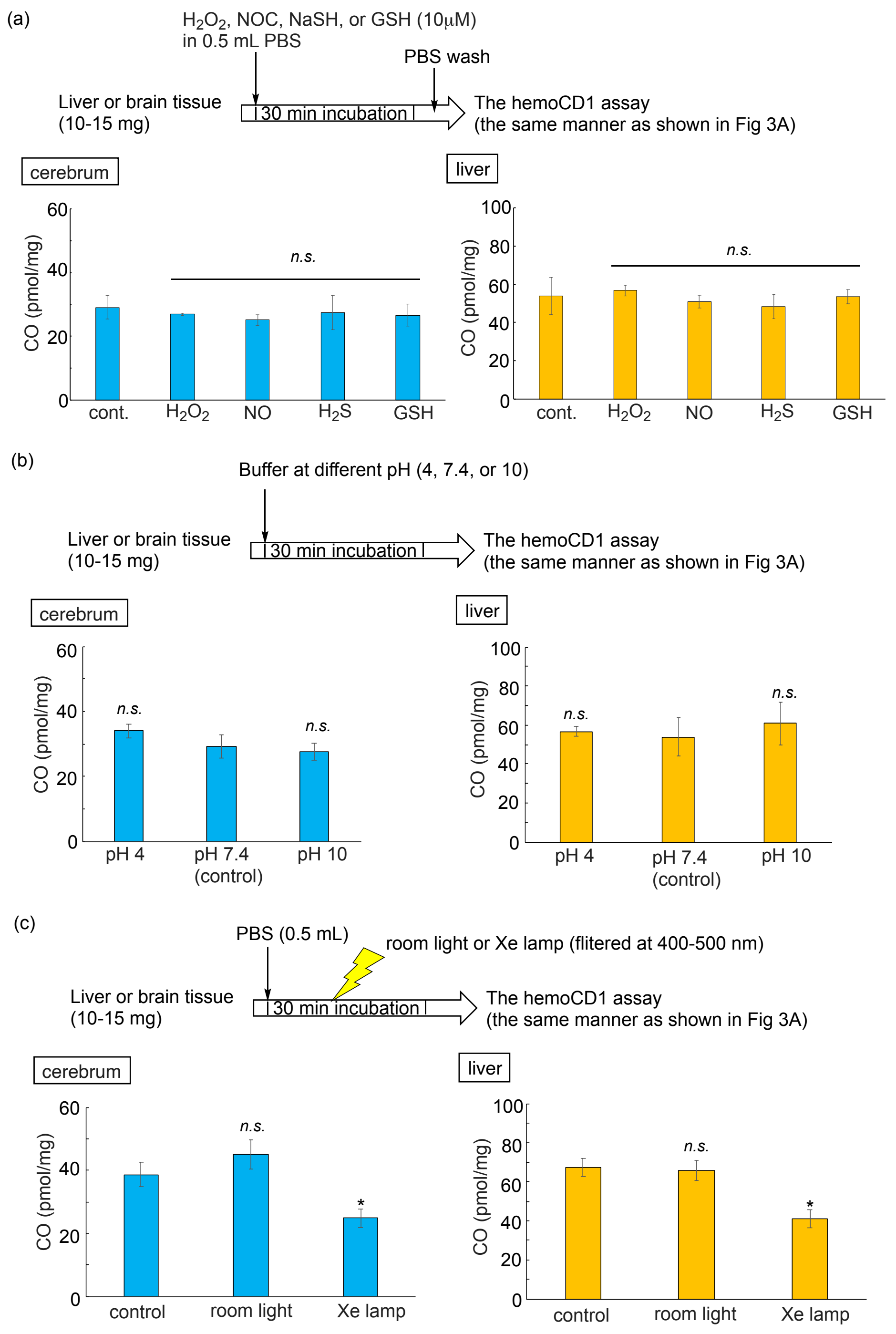


**Fig. S11. Effects of external reactive species (a), pH (b), and light (c) on quantification of endogenous CO in the liver and brain tissues by hemoCD1.** The data indicate that the assay using hemoCD1 was unaffected by the reactive species, $H_2O_2$, NO, $H_2S$ and GSH, pH (from 4 to 10), and room light. The data in c indicate that the strong irradiation by Xe lamp reduced CO due to dissociation of the CO-Fe complex. Statistical significance, $^*p < 0.05$ vs control; not significant, *n.s.*

a)

b)

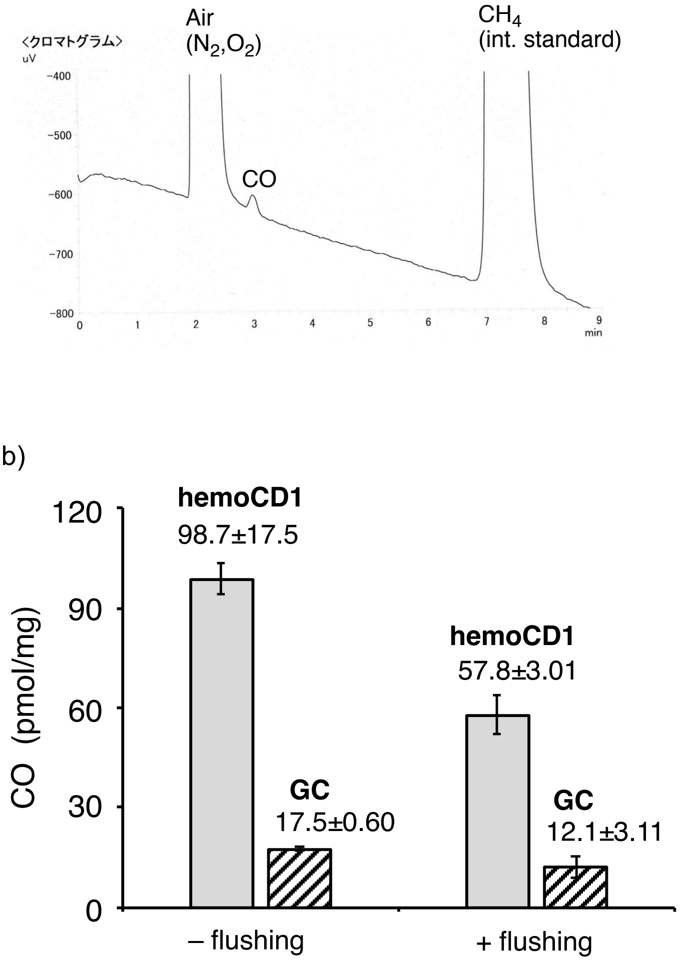


**Fig. S12**. **Comparison between the hemoCD1 assay and the gas chromatography (GC) method for CO quantification in tissues.** (a) A typical gas chromatogram recording the amount of CO from a liver sample. The headspace gas was analyzed by a TCD detector as reported in the literature, (see Ref 16 in the text). (b) The amounts of CO in the liver tissues (– or + flushing) quantified by the hemoCD1 assay and the GC method. Each bar represents mean ± SE (*n* = 3–6).

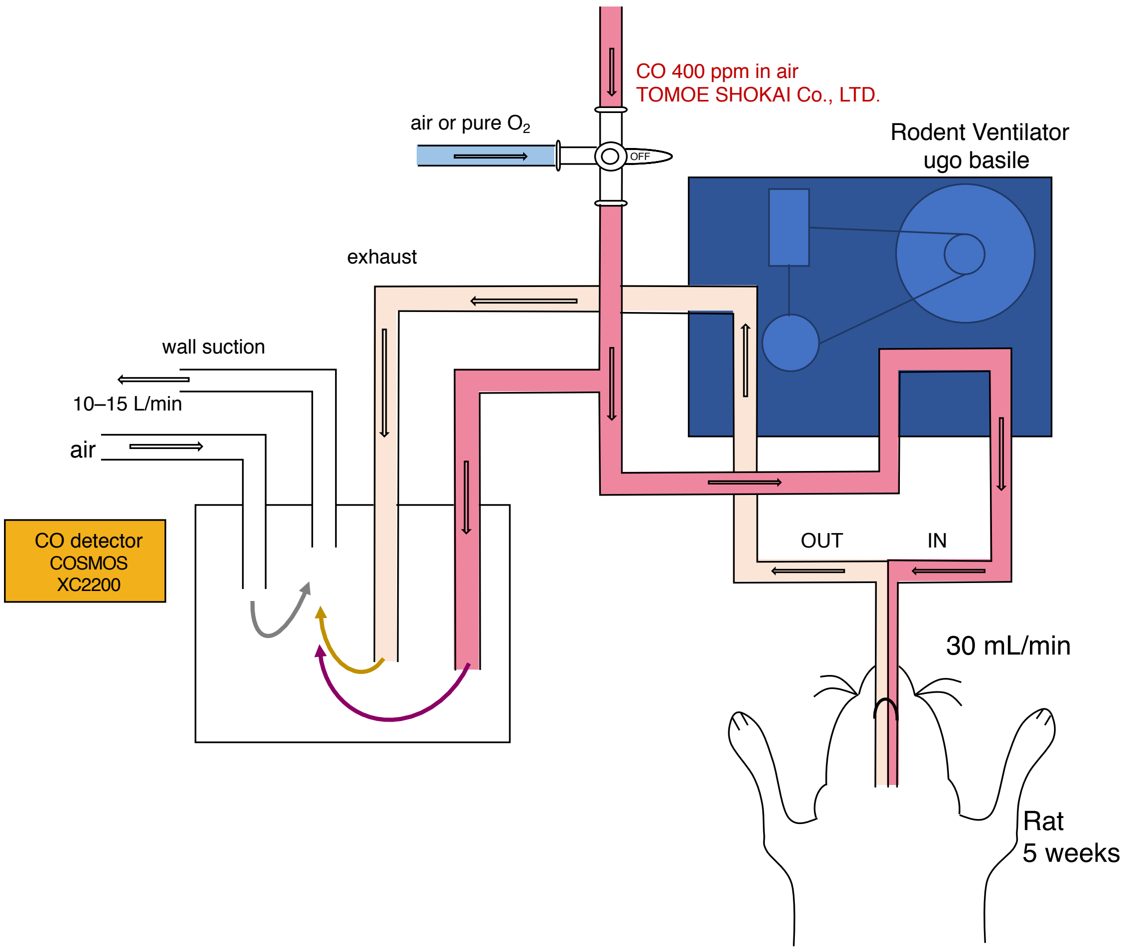


**Fig. S13. Schematic representation of the experimental setup for CO gas inhalation in rats.** The apparatus allowed a rapid switch from inhaled CO gas (400 ppm) to air or $O_2$ by handling the three-way stop cock. The rate of inhalation was controlled by the rodent ventilator. The exhaled gas was mixed with air before exiting. For safety reasons, a CO detector was used for monitoring the atmospheric concentrations of CO.

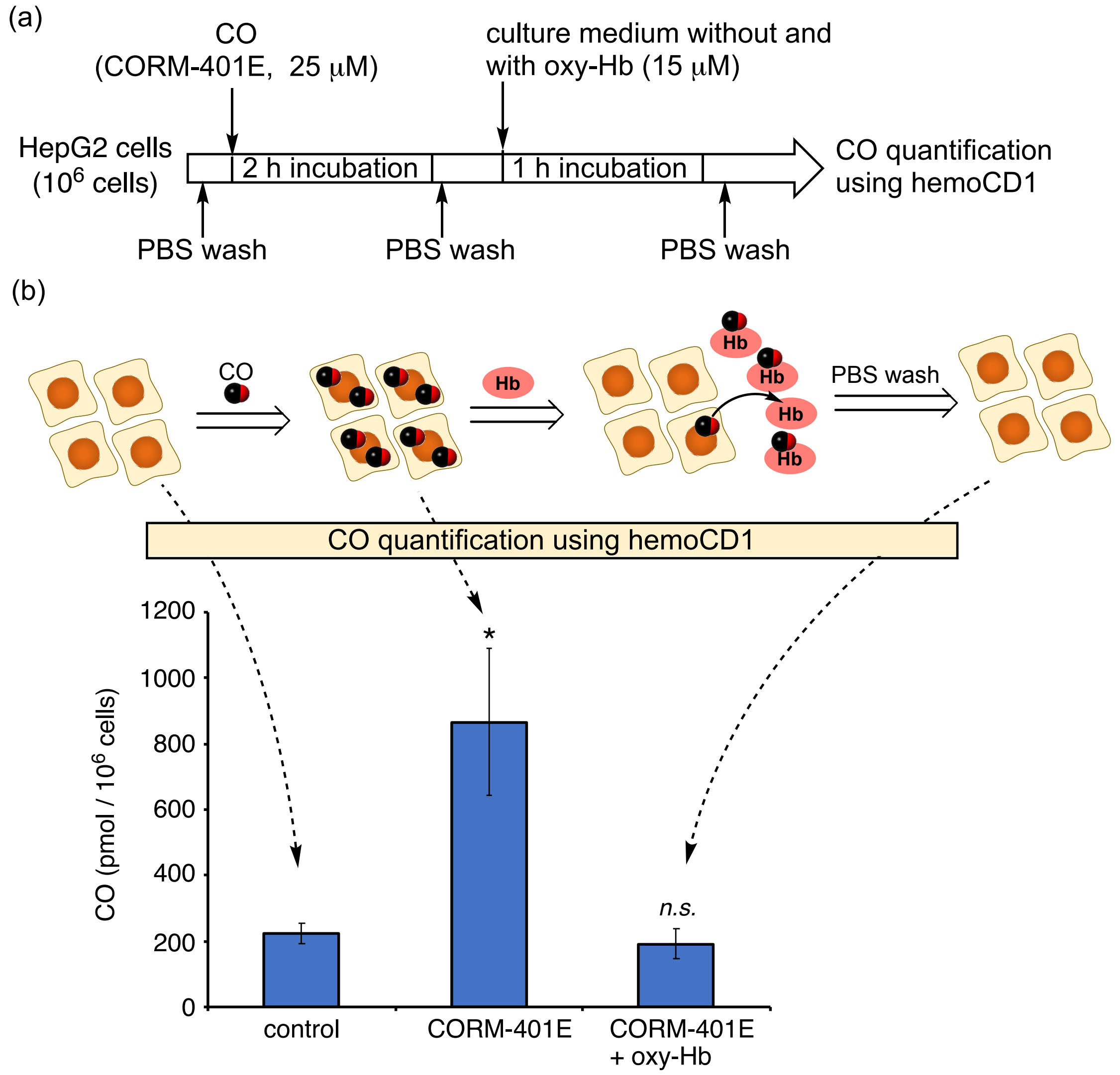


**Fig. S14. *In vitro* experiments to demonstrate CO transfer from tissues to Hb.** (a) Hepatocytes (1 x $10^6$) were treated for 2 h with 25 μM CORM-401E, a CO-releasing molecule recently synthesized in our laboratory for efficient intracellular CO delivery tool (unpublished data, patent pending: PCT/JP2020/006033). Cells were washed out and then treated with medium in the presence of oxy-Hb. Finally, after removing the medium containing Hb, the amount of residual CO in cells was quantified by the hemoCD1 assay. (b) The amounts of CO in cells determined by the assay using hemoCD1 at each step. The results show that cells incubated with CORM401-E stored CO inside, which was then transferred when incubated with oxy-Hb. Each bar represents mean ± SE ($n$ = 3). Statistical significance, *$p < 0.05$; not significant, *n.s.* versus control.

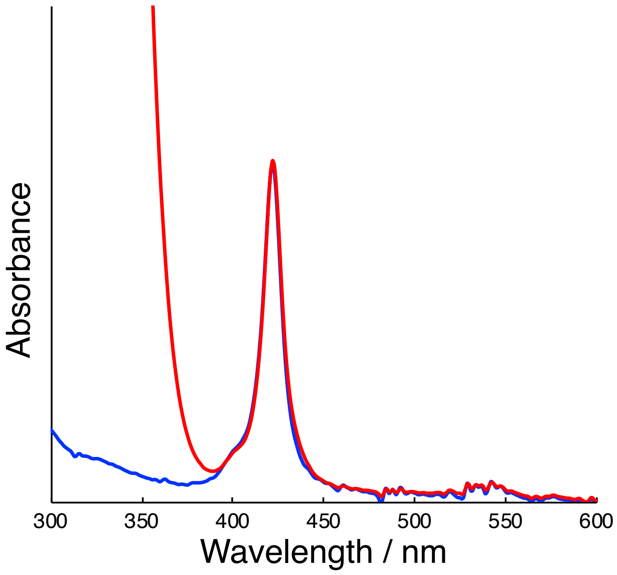


**Fig. S15**. **UV-vis spectra of rat urine collected after i.v. infusion of oxy-hemoCD1 to CO-treated rats before (blue) and after the addition of $Na_2S_2O_4$ (red).** The completely overlapped spectral shapes indicates that hemoCD1 in the urine was almost 100% in the CO-bound form. A similar spectral profile has been previously reported by us (Refs 35,36).

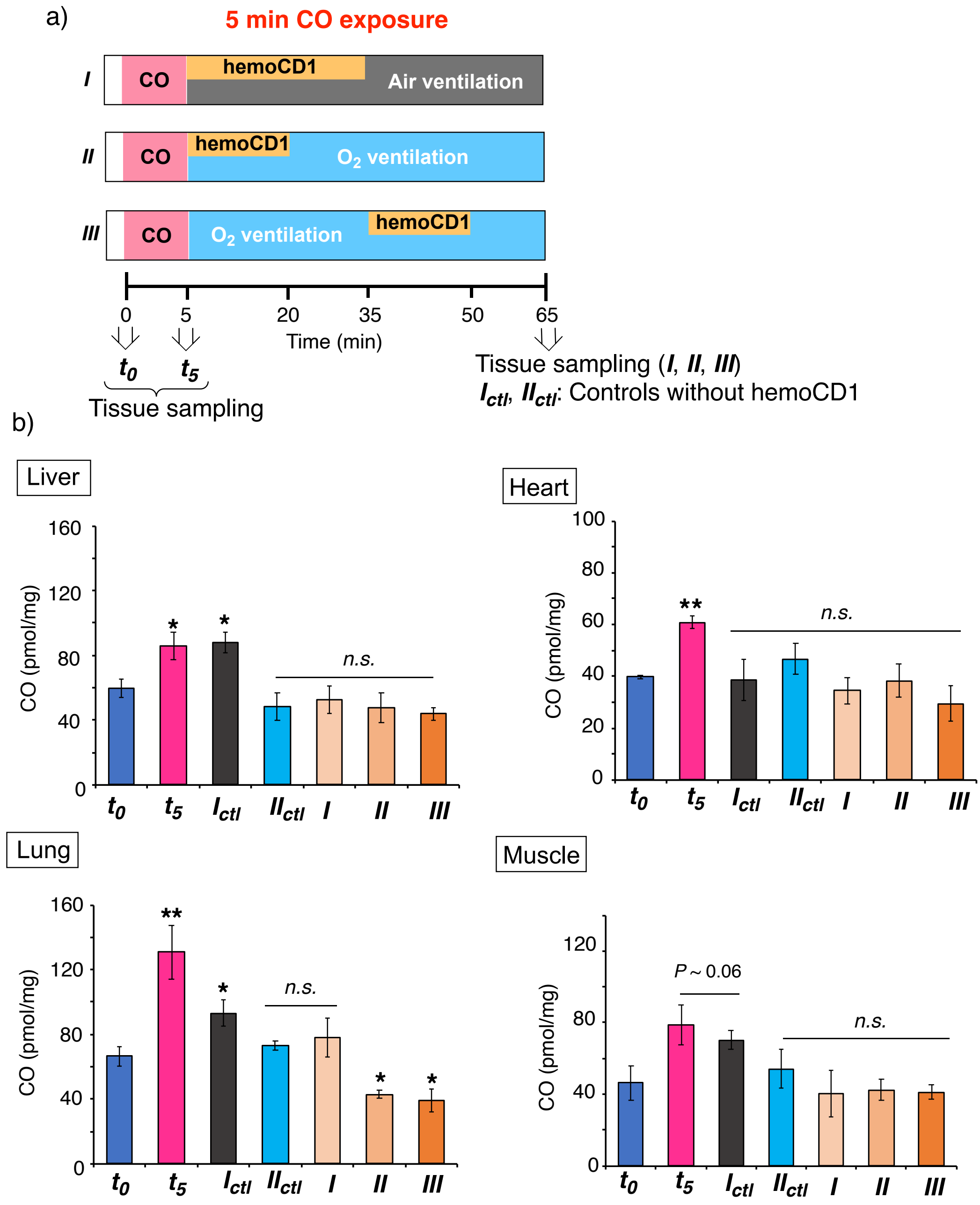


**Fig. S16. Effect of air/$O_2$ ventilation in combination with hemoCD1 on CO distribution in tissues after CO inhalation *in vivo*.** (a) Experimental protocols used are the same as those in Fig. 6A. (b) Amounts of CO detected in liver, heart, lung and muscle tissues collected as indicated in (a). Each bar represents the mean ± SE ($n$ = 3–6). Statistical significance, *$p < 0.05$, **$p < 0.01$; not significant, *n.s.* versus $\boldsymbol{t_0}$.

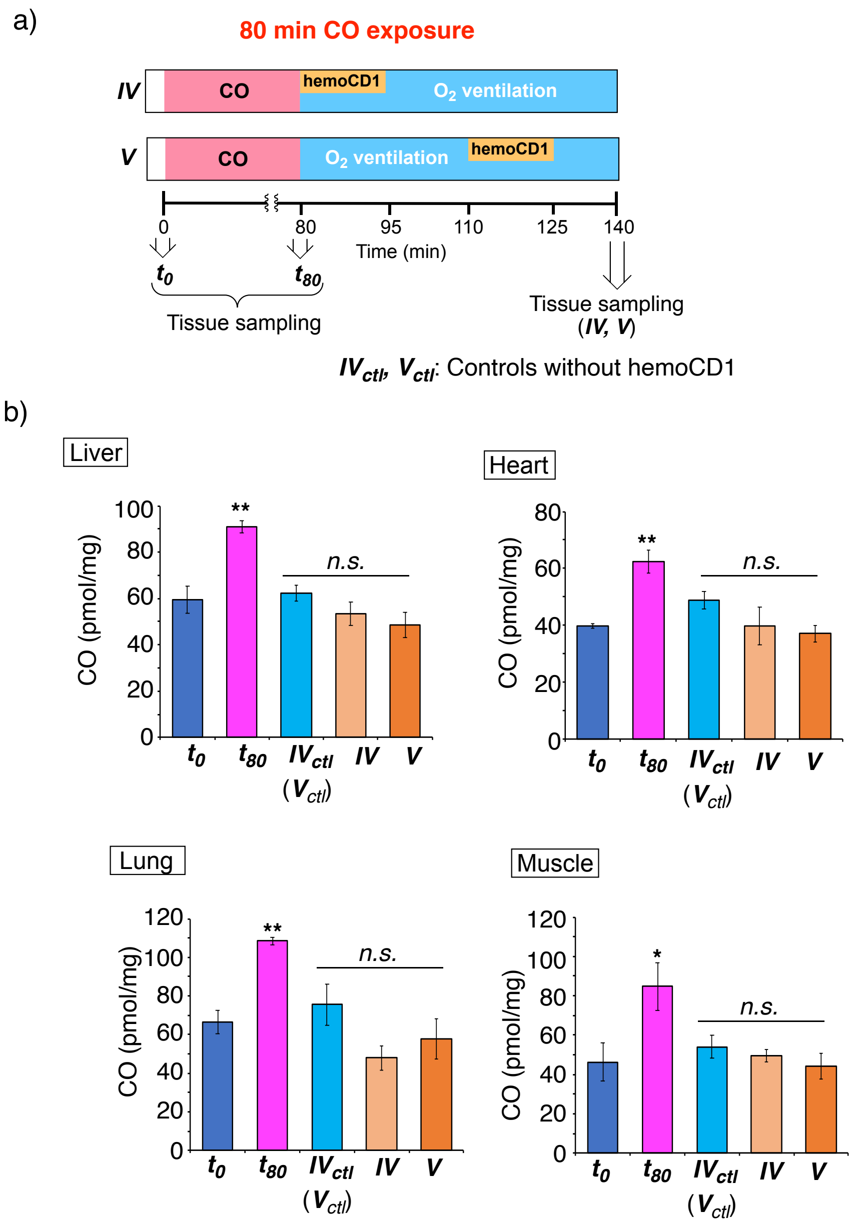


**Fig. S17**. **Effect of $O_2$ ventilation in combination with hemoCD1 on CO distribution in tissues after CO inhalation *in vivo*.** Experimental protocols used are the same as those in Fig. 6B. (b) Amounts of CO detected in liver, heart, lung and muscle tissues collected as indicated in (a). Each bar represents the mean ± SE ($n$ = 4–6). Statistical significance, $^*p < 0.05$, $^{**}p < 0.01$; not significant, *n.s.* versus $\boldsymbol{t_0}$.
